# LSD1 ablation promotes mammary tumor metastasis by attenuating NK cell-mediated anti-tumor immunity

**DOI:** 10.64898/2026.03.12.711410

**Authors:** Dongxi Xiang, Sen Han, Aina He, Guyu Qin, Roderick T. Bronson, Zhe Li

**Affiliations:** Division of Genetics, Brigham and Women’s Hospital, Boston, Massachusetts 02115, USA; Department of Medicine, Harvard Medical School, Boston, Massachusetts 02115, USA; Department of Urology, Boston Children’s Hospital, Boston, Massachusetts 02115, USA; Rodent Histopathology, Harvard Medical School, Boston, Massachusetts 02115, USA

**Keywords:** Epigenetic regulator, LSD1 (KDM1A), luminal breast cancer metastasis, immune microenvironment, NK cells

## Abstract

Epigenetic deregulation can alter the expression of cancer-related genes in tumor cells and may promote metastasis by influencing interactions between tumor cells and their immune microenvironment. However, the underlying immune mechanisms remain poorly understood. LSD1 (KDM1A) is a histone demethylase that has been proposed to function as a tumor and metastasis suppressor in breast cancer. Here, using the *MMTV-PyMT* breast cancer mouse model, we show that natural killer (NK) cells play a critical role in suppressing tumor cell metastasis to the lung, and that ablation of LSD1 leads to increased lung metastasis. This phenotype is accompanied by pronounced upregulation of immune-related genes, including major histocompatibility complex class I (MHC-I) genes, in tumor cells and by extensive remodeling of the tumor immune microenvironment, characterized by reduced abundance and maturation of NK cells. Consistent with these observations, NK cells exhibit reduced cytotoxicity toward *Lsd1*-null *PyMT* tumor cells. Notably, NK cell-mediated killing can be restored by disrupting expression of the non-classical MHC-I molecule Qa-1, a ligand for the inhibitory NK receptor CD94/NKG2A, in tumor cells. In transplantation experiments, *Lsd1*-null *PyMT* tumor cells formed significantly larger lung metastatic lesions than *Lsd1*-wildtype tumor cells in SCID mice, which possess functional NK cells, but not in NSG mice that lack NK cells. Collectively, these findings suggest that epigenetic deregulation in LSD1-deficient mammary tumor cells reprograms the tumor immune microenvironment, resulting in impaired NK cell-mediated tumor surveillance and enhanced metastatic progression.

## Introduction

Most cancer-related deaths are caused by metastasis ^1, 2^. Large-scale sequencing studies have identified the majority of driver mutations responsible for the transformation of normal cells into cancer cells ^3^. However, similar efforts to identify genetic mutations that specifically drive metastasis have been largely unsuccessful ^3, 4^, leading to the notion that metastasis may not be driven by distinct genetic alterations ^3^. Genetic mutations predicted to be drivers in metastatic lesions are typically already present in the corresponding primary tumors, although their frequencies may be higher in metastases ^2, 5–7^. The lack of metastasis-specific genetic drivers has therefore led to the hypothesis that epigenetic mechanisms, through which cancer cells acquire pro-metastatic transcriptional programs, may play a central role in metastatic progression ^1, 2^.

Epigenetic regulatory machinery also plays an essential role in safeguarding cells from aberrant immune responses, for example by repressing endogenous retroviruses (ERVs) within the genome ^8–10^. Epigenetic deregulation can unleash ERV expression, which in turn can trigger innate immune responses. Sustained activation of these pathways may result in chronic inflammation and the establishment of an immunosuppressive microenvironment that promotes tumor progression ^8–10^. However, how immune mechanisms triggered by epigenetic dysregulation contribute to metastatic progression remains largely unclear.

The epigenetic landscape of a cell is shaped by diverse epigenetic regulators. Lysine-specific demethylase 1 (LSD1, also known as KDM1A) is a key histone demethylase that can function as a transcriptional repressor by removing methyl groups from mono- and di-methylated lysine 4 on histone H3 (H3K4me1/me2) ^11–13^. LSD1 can also act as a transcriptional activator through demethylation of mono- and di-methylated lysine 9 on histone H3 (H3K9me1/me2) ^14–16^. In breast cancer, LSD1 has been reported to function either as a tumor suppressor ^13, 17, 18^ or as an oncogenic factor ^19–23^. These apparently contradictory roles may reflect the context-dependent functions of LSD1 in different mammary epithelial cell (MEC) populations. Alternatively, they may arise from distinct roles of LSD1 in shaping the tumor microenvironment, particularly the immune microenvironment.

The immune system plays a critical role in dictating the metastatic spread of breast cancer. On one hand, immune effector cells such as CD8⁺ T cells and NK cells can recognize and eliminate tumor cells, thereby restricting metastatic outgrowth. On the other hand, tumor cells can recruit immunosuppressive cell populations or promote the differentiation of tumor-infiltrating immune cells toward pro-tumorigenic states that facilitate metastasis ^24^. The recent clinical success of immune checkpoint blockade (ICB) therapy has generated strong interest in immunotherapy for the treatment of multiple cancers, including breast cancer ^25^. However, the response rate to ICB therapy remains relatively low ^26, 27^. To improve therapeutic efficacy, ongoing efforts focus on combining ICB with other treatment modalities ^25^, including epigenetic therapies ^28, 29^. This strategy is based on the observation that epigenetic drugs, such as inhibitors of DNA methyltransferases (DNMTs) or Histone deacetylases (HDACs), can enhance anti-tumor immune responses by modulating key pathways involved in tumor-immune interactions ^28, 29^. More recently, inhibition of LSD1 has been shown to induce a type I interferon (IFN) response, enhance tumor immunogenicity, stimulate anti-tumor T cell immunity, and improve responses to anti-PD-(L)1 immunotherapy ^30^.

Despite these promising findings, LSD1 disruption has also been linked to increased breast cancer metastasis in several studies ^13, 18, 31^. How LSD1 inhibition-based epigenetic therapy influences metastatic progression therefore remains unclear. In this study, we demonstrate that disruption of LSD1 in mammary tumor cells promotes lung metastasis, at least in part through attenuation of NK cell-mediated anti-tumor immunity.

## Results

### Induced loss of LSD1 in *PyMT* tumor cells increases metastasis

We reported previously that in luminal breast cancer, LSD1 largely functions as a tumor/metastasis suppressor, in part by controlling MEC-intrinsic transcriptional programs that restrain invasion and migration of luminal MECs ^18^. Our experimental approach utilized intraductal injection of a Cre-expressing adenovirus driven by the Keratin 8 promoter (*Ad-K8-Cre*) ^32, 33^ into mammary glands (MGs) of the autochthonous luminal breast cancer mouse model *MMTV-PyMT* (hereafter referred to as *PyMT*) ^34^. The *PyMT* mice also carried *Lsd1* conditional knockout (KO) alleles (*Lsd1^L/L^*) together with a conditional *Rosa26-LSL-YFP* (*R26Y*) Cre-reporter ^35^. Intraductal injection of *Ad-K8-Cre* resulted in inactivation of *Lsd1* while simultaneously activating YFP expression in the same *PyMT* tumor cells (Figure 1A). Using this approach, we observed significant increases in both the number and size of lung metastatic lesions in *PyMT* mice with induced loss of LSD1 compared with *PyMT* mice retaining wild-type (wt) LSD1 (Figure 1B).

**Figure 1.**
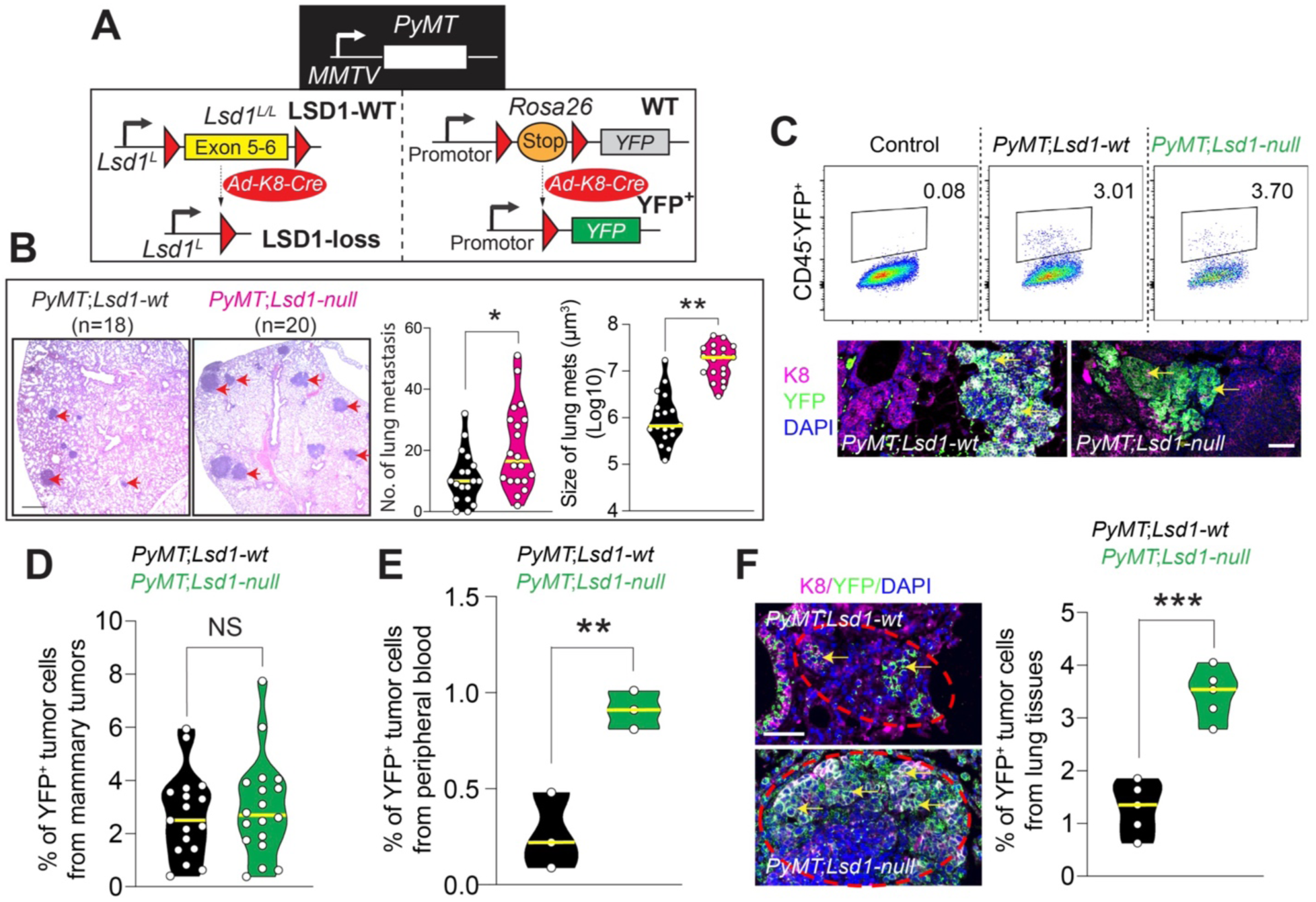
Induced loss of LSD1 in *PyMT* mammary luminal cells led to increased lung metastasis. **(A)** Schematic diagram showing disruption of *Lsd1* and activation of the YFP reporter in mammary luminal cells induced by intraductal injection of *Ad-K8-Cre*. **(B)** Left: Hematoxylin and Eosin (H&E) images showing representative lung metastatic lesions (arrows) of mice with the indicated genotypes; right: numbers and sizes of metastatic lesions as indicated in the left panel. mets: metastases. Scale bar: 500 µm. **(C)** Top: representative FACS gating of CD45⁻YFP^+^ tumor cells in *PyMT* mice with/without induced LSD1-loss; bottom: co-immunofluorescence (co-IF) staining showing YFP^+^ tumor lesions (arrows). Scale bar: 50 µm. **(D)** Quantification of % of YFP^+^ cells in primary mammary tumors in mice as indicated in (C) top panel. **(E)** Quantification of % of YFP^+^ tumor cells from peripheral blood in mice as indicated. **(F)** Left: co-IF staining showing YFP^+^ lung metastatic lesions (arrow) in *PyMT* female mice with/without induced LSD1-loss; right: quantification of % of YFP^+^ cells in lungs of mice as indicated in the left panel. *P* value: *p<0.05, **p<0.01, ***p<0.005, NS = not significant, two-tailed Student’s *t*-test. Data represent mean ± SEM. Yellow lines in violin plots represent mean values.

Since intraductal injection of *Ad-K8-Cre* induced *Lsd1* deletion in only a subset of *PyMT* tumor cells, this approach generated a mosaic system in the injected *MMTV-PyMT;Lsd1^L/L^;R26Y* females. In this system, YFP⁺ tumor cells lack LSD1 expression, whereas YFP⁻ tumor cells retain LSD1 expression. In contrast, in similarly injected *MMTV-PyMT;R26Y* control females, both YFP⁺ and YFP⁻ tumor cells remain positive for LSD1 expression. We therefore analyzed YFP⁺ cells in tumors from both experimental and control *PyMT* females. In primary mammary tumors, the two groups showed similar percentages of CD45⁻YFP⁺ tumor cells (Figure 1C-D and Figure S1A). In peripheral blood, however, the proportion of circulating CD45⁻YFP⁺ tumor cells in *PyMT* mice with induced LSD1-loss was ∼3.5-fold higher than in control *PyMT* females with wt LSD1 (Figure 1E and Figure S1B). In lung metastatic lesions, those from injected *MMTV-PyMT;Lsd1^L/L^;R26Y* experimental females contained on average ∼2.7-fold higher percentages of YFP^+^ cells than lesions from control females (Figure 1F). Together, these data indicate that although *Lsd1*-null *PyMT* tumor cells do not exhibit a pronounced clonal growth advantage in primary mammary tumors, they possess a significant advantage in seeding the lung and forming metastatic lesions.

### LSD1-loss in *PyMT* tumor cells alters immune-related gene expression

The increased metastatic capacity of *Lsd1*-null *PyMT* tumor cells may arise from both cell-intrinsic and extrinsic mechanisms ^18^. To gain further insights into the enhanced metastatic potential of *Lsd1*-null *PyMT* tumor cells, we sorted YFP^+^ cells from primary tumors arising in the MGs of *MMTV-PyMT;Lsd1^L/L^;R26Y* females (i.e., YFP⁺ *Lsd1*-null *PyMT* tumor cells) and compared them with YFP^+^ *Lsd1*-wt *PyMT* tumor cells sorted from *MMTV-PyMT;R26Y* control females ∼2-3 months after intraductal injection of *Ad-K8-Cre*. These cells were subjected to RNA-sequencing (RNA-seq).

Strikingly, gene set enrichment analysis (GSEA) ^36^ of the resulting expression profiles revealed that many of the top enriched gene sets in *Lsd1*-null *PyMT* tumor cells were associated with immune response, including pathways related to cytotoxic T cells and immune checkpoint regulation (Figure 2A and Figure S2A-C). Among immune-related genes, we noted that most major histocompatibility complex class I (MHC-I) genes (both classical and non-classical), as well as genes involved in immune checkpoint pathways (Figure 2B) and IFN signaling (Figure 2C), were upregulated in *Lsd1*-null *PyMT* tumor cells. To validate these findings, we utilized YFP^+^ *Lsd1*-null and *Lsd1*-wt *PyMT* cells sorted from MGs of *PyMT* mice ∼2-weeks after intraductal injection of *Ad-K8-Cre* and maintained *ex vivo* as organoids. In parallel, we generated CRISPR-mediated KO of *Lsd1* in *PyMT* tumor organoid cells. In both organoid systems, quantitative RT-PCR (qRT-PCR) confirmed upregulation of immune-related genes in *Lsd1*-null *PyMT* organoid cells (Figure 2D and Figure S2D). In addition, we treated *PyMT* tumor organoids with the LSD1 inhibitor GSK2879552 or HCI-2509 and measured expression levels of several key immune genes by qRT-PCR. We found that pharmacological inhibition of LSD1 in *PyMT* tumor cells also upregulated expression of these immune genes in them (Figure S2E).

**Figure 2.**
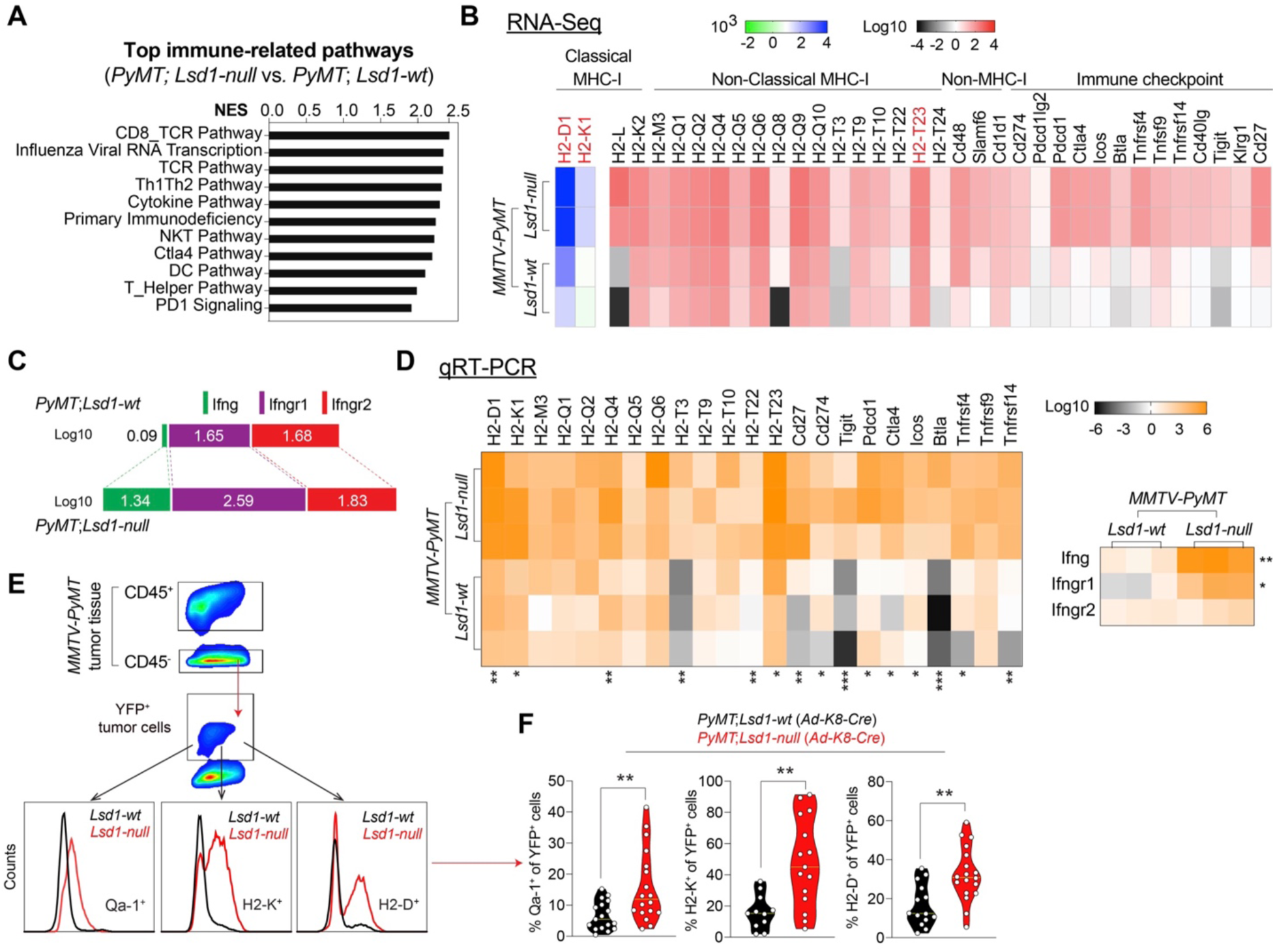
LSD1-loss led to upregulation of immune-related genes in *PyMT* tumor cells. **(A**) The majority of top-enriched gene sets in sorted YFP^+^ *Lsd1*-null (vs. *Lsd1*-wt) *PyMT* tumor cells are immune-related. NES: normalized enrichment score. **(B**) Heatmaps showing upregulation of most classical and non-classical MHC-I genes as well as select non-MHC-I genes in sorted YFP^+^ *Lsd1*-null (vs. *Lsd1*-wt) *PyMT* tumor cells. **(C)** Increased expression of IFNγ (*Ifng*) and its receptor genes in *Lsd1*-null (vs. *Lsd1*-wt) *PyMT* tumor cells. Data in (B-C) are based on RNA-seq. **(D)** qRT-PCR validation of selected genes in (B) and (C) in tumor organoid cells. **(E)** Representative FACS plots showing increased levels of MHC-I molecules (Qa-1, H2-K, and H2-D) in CD45^-^YFP^+^ *PyMT* tumor cells with LSD1-loss. **(F)** Quantification of % of CD45⁻YFP^+^ cells positive for the indicated MHC-I molecules as in (E). *P* value: *p<0.05, **p<0.01, ***p<0.005, ****p<0.001, two-tailed Student’s *t*-test. Data represent mean ± SEM.

In CRISPR-induced *Lsd1*-KO *PyMT* cells, as well as in *PyMT* tumor organoids treated with the LSD1 inhibitor GSK2879552 or HCI-2509, we also observed upregulation of transcripts derived from several mouse ERVs ^37, 38^ (Figure S2F-G). This likely reflects derepression of ERV loci following loss of LSD1-mediated transcriptional silencing ^30^. Together, these findings suggest that the observed changes in immune-related gene expression in *Lsd1*-null *PyMT* cells (i) are unlikely to result from contamination of immune cells (e.g., macrophages) within the sorted YFP^+^ population and (ii) instead largely represent an early cell-intrinsic response in *PyMT* MECs following LSD1-loss. One possible mechanism is activation of innate immune signaling in response to double-stranded RNA stress caused by ERV derepression, similar to findings in human breast cancer cells with LSD1 ablation ^30^.

Lastly, to validate the upregulation of MHC-I genes at the protein level, we performed fluorescence-activated cell sorting (FACS) analysis for the MHC-I molecules H2-D and H2-K, as well as the non-classical MHC-I molecule Qa-1 (encoded by *H2-T23*) (Figure S3A), in *PyMT* tumors with or without induced LSD1-loss. We observed higher fluorescence intensities for all three MHC-I molecules, along with significant increases in the proportions of H2-D^+^, H2-K^+^ and Qa-1^+^ cells among CD45⁻YFP⁺ tumor cells lacking LSD1 (Figure 2E-F).

### LSD1-loss in *PyMT* tumor cells reprograms the immune microenvironment

The profound changes in the expression of immune-related genes in *Lsd1*-null *PyMT* tumor cells suggested that these alterations might reshape the tumor immune microenvironment, potentially contributing to the increased metastatic phenotype through MEC-extrinsic mechanisms. To test this possibility, we performed multi-color FACS analysis (Figure S3A) of primary *PyMT* tumors arising in the MGs, as well as peripheral blood samples from these mice. In primary tumors, we observed a significant increase in the CD45^+^ leukocyte population in tumors with induced LSD1-loss (Figure 3A). Among CD45^+^ cells, tumors with LSD1-loss exhibited a significant increase in the CD4⁺ T cell population, accompanied by reduced frequencies of CD8⁺ T cells, NK cells, and macrophages (Figure 3B and Figure S3B-C). Since *PyMT* tumors with LSD1-loss contained a larger overall CD45⁺ leukocyte compartment, we also calculated the proportions of individual immune populations among total live cells within the tumors. This analysis similarly revealed increased CD4⁺ T cells and decreased NK cells and macrophages in tumors with induced LSD1-loss (Figure 3B). Together, these results indicate that induced loss of LSD1 in *PyMT* tumor cells leads to substantial reprogramming of the tumor immune microenvironment.

**Figure 3.**
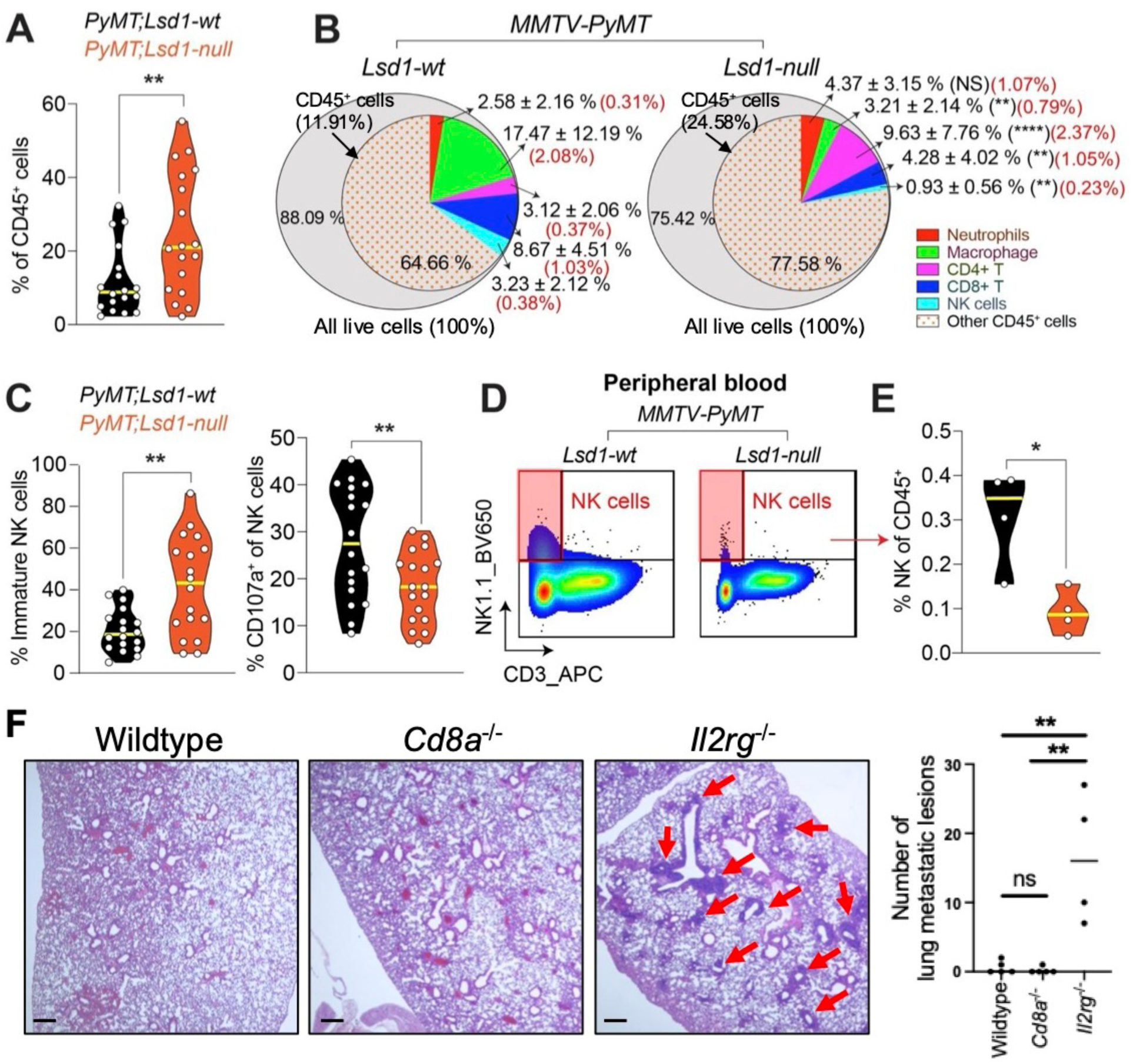
LSD1-loss in *PyMT* luminal cells reprogrammed the tumor immune microenvironment. **(A)** Alteration of CD45^+^ cells in *PyMT* mammary tumors 2-3 months after induced LSD1-loss. **(B)** Pie charts summarizing the mean abundance [shown as % within CD45^+^ cells (=100%, inner cycle); the statistical significance was calculated based on this %] of immune cell subsets in *PyMT* tumors with or without LSD1-loss; mean % of each immune cell population within live cells (100%, outer circle) is provided in the parentheses in red font. **(C)** Alteration of the immature NK cell subset and activated CD107a^+^ NK cells in *PyMT* tumors with LSD1-loss (*Lsd1-null*). **(D)** Representative FACS plots showing decreased % of NK cells in peripheral blood of *PyMT* female mice with induced LSD1-loss. **(E)** Quantification of % of NK cells as indicated in (D). **(F)** *PyMT* tumor organoids were inoculated into C57BL/6 syngeneic recipient female mice with the indicated genotypes via intraductal injection; H&E images show representative lung metastatic lesions (red arrows, left panel), summarized data for the number of lung metastatic lesions is shown in the right panel. Scale bar: 500 μm. *P* value: *p<0.05, **p<0.01, ***p<0.005, ****p<0.001, ns = not significant, two-tailed Student’s *t*-test. Data represent mean ± SEM.

The strong upregulation of MHC-I genes in *Lsd1*-null *PyMT* tumor cells (Figure 2B and D), together with the reduced NK cell population observed in primary *PyMT* tumors with induced LSD1-loss (Figure 3B and Figure S3B-C), raised the possibility that NK cells may contribute to the differences in lung metastatic burden between *PyMT* mice with or without LSD1-loss. We therefore examined NK cells in greater detail. NK cell maturation in primary tumors was assessed by FACS staining for CD11b and CD27, markers commonly used to define NK cell maturation stages [NK cell maturation is determined as: CD11b^low^ CD27^low^ (most immature) -> CD11b^low^ CD27^high^ -> CD11b^high^ CD27^high^ -> CD11b^high^ CD27^low^ (most mature) ^39^]. In addition, we analyzed CD107a, a degranulation marker associated with NK cell activation ^40^ (Figure S3B).

In *PyMT* tumors with induced LSD1-loss, we observed not only a reduction in the overall NK cell population but also a significant increase in the immature CD11b^low^ CD27^low^ NK cell subset, as well as a pronounced reduction in the mature CD107a^+^ NK cell subset (Figure 3C). Importantly, in peripheral blood, the frequency of the NK cells in *PyMT* mice with induced LSD1-loss was also lower than that in control *PyMT* females (Figure 3D-E and Figure S1B).

To directly assess the relative contributions of NK cells versus other immune populations (e.g., CD8⁺ T cells) in restricting lung metastasis of *PyMT* tumor cells, we transplanted equal numbers of *PyMT* tumor organoid cells into syngeneic WT, *Cd8a^-/-^*, and *Il2rg^-/-^* female recipient mice. Of note, *Cd8a^-/-^* and *Il2rg^-/-^* mice lack CD8^+^ T cells and NK cells ^41, 42^, respectively. We found that lungs from *Il2rg^-/-^* recipients developed significantly more metastatic lesions than those from either *Cd8a^-/-^* or WT recipients (Figure 3F). These results support a critical role for NK cells in suppressing *PyMT* tumor cell metastasis to the lung. Collectively, our data suggest that induced LSD1-loss in the *PyMT* model leads to both reduced abundance and impaired maturation of NK cells, which may contribute to the enhanced lung metastasis observed in this setting.

### Induction of LSD1-loss in *PyMT* tumor cells using a CreER/tamoxifen system produces a similar phenotype

Intraductal injection of adenovirus can trigger immune response in the injected MG, although this response typically resolves over time ^32^. To exclude the possibility that the adenoviral injection procedure itself contributed to the increased lung metastasis phenotype, we generated *MMTV-PyMT;K8-CreER;Lsd1^L/L^;R26Y* females and inactivated *Lsd1* in the same K8^+^ *PyMT* tumor cells through tamoxifen injection, which induces CreER activity and Cre-mediated disruption of *Lsd1* together with activation of YFP expression (Figure 4A). Of note, we reported previously that this CreER/tamoxifen-based approach targets K8^+^ luminal MECs at a frequency comparable to that achieved with the *Ad-K8-Cre* intraductal injection approach ^33^.

**Figure 4.**
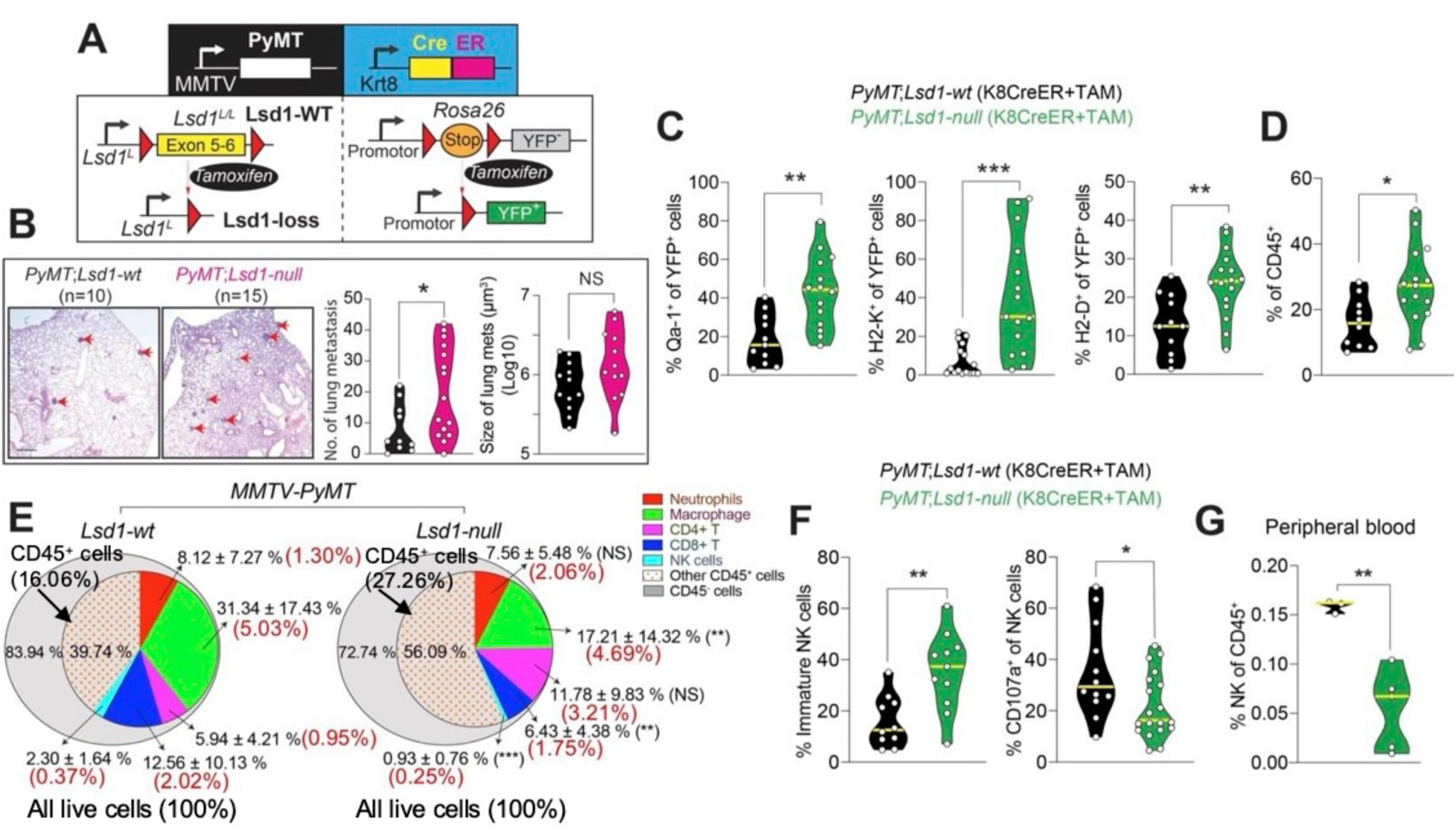
CreER/tamoxifen-induced LSD1-loss in the *PyMT* model led to increased lung metastasis and reprogrammed tumor immune microenvironment. **(A**) Schematic diagram showing disruption of *Lsd1* and activation of the YFP reporter in *MMTV-PyMT;K8-CreER;Lsd1^L/L^;R26Y* female mice induced by tamoxifen injection (TAM). **(B**) Left: H&E images showing representative lung metastatic lesions (arrows) of mice with the indicated genotypes; right: numbers and sizes of lung metastatic lesions as indicated in the left panel. mets: metastases. Scale bar: 500 µm. **(C)** Quantification of % of cells positive for MHC-I molecules (Qa-1, H2-K, and H2-D) in CD45⁻YFP^+^ *PyMT* tumor cells with or without TAM-induced LSD1-loss. **(D)** Alteration of CD45^+^ cells in *PyMT* tumors with TAM-induced LSD1-loss. **(E)** Pie charts summarizing the mean abundance [shown as % within CD45^+^ cells (=100%, inner cycle); the statistical significance was calculated based on this %] of immune cell subsets in *PyMT* tumors with or without TAM-induced LSD1-loss; mean % of each immune cell population within live cells (100%, outer circle) is provided in the parentheses in red font. **(F)** Alteration of the immature NK cell subset and activated CD107a^+^ NK cells in *PyMT* tumors with TAM-induced LSD1-loss. **(G)** Decreased % of NK cells in peripheral blood of *PyMT* female mice with TAM-induced LSD1-loss. Sample legends in (D) and (G) are the same as in (C). *P* value: *p<0.05, **p<0.01, ***p<0.005, NS = not significant, two-tailed Student’s *t*-test. Data represent mean ± SEM.

Using this second genetic strategy, we observed similar increases in both the number and, to a lesser extent, the size of lung metastatic lesions in *PyMT* mice with induced LSD1-loss (Figure 4B). FACS analysis revealed significantly increased expression of the MHC-I molecules H2-D, H2-K, and Qa-1, in CD45⁻YFP⁺ *PyMT* tumor cells from MGs with induced LSD1-loss (Figure 4C). Within the tumor microenvironment of primary *PyMT* mammary cancers with induced LSD1-loss, we also observed an increase in the total CD45^+^ leukocyte population (Figure 4D). Among CD45^+^ leukocytes, significant decreases were detected in the CD8⁺ T cell, NK cell, and macrophage subsets (Figure 4E and Figure S4A-B). When normalized to the percentage of total live cells, *PyMT* tumors with induced LSD1-loss displayed reduced NK cell, CD8⁺ T cell, and macrophage populations, along with an increased CD4⁺ T cell population (Figure 4E). Among tumor-associated NK cells, we further observed a significant increase in the immature NK cell subset accompanied by a reduction in the mature NK cell subset (Figure 4F). Consistent with these findings, the frequency of NK cells in the peripheral blood of *PyMT* mice with tamoxifen-induced loss of LSD1 was also lower than that observed in control *PyMT* females (Figure 4G). Collectively, these results indicate that it is the induced loss of LSD1 in *PyMT* tumor cells that drives the observed reprogramming of the tumor immune microenvironment and the enhanced metastatic phenotype in the *PyMT* model.

### LSD1-loss in *PyMT* tumor cells attenuates NK cell-mediated anti-tumor immunity

NK cells are innate lymphocytes that play a critical role in controlling metastatic dissemination (^43^ and Figure 3F). NK cell activity is regulated through the integration of signals from activating and inhibitory receptors ^44^. In addition to their role in antigen presentation to CD8^+^ T cells, MHC-I molecules can bind inhibitory receptors on NK cells and suppress their activation, thereby preventing the killing of healthy “self” cells ^45, 46^. The increased MHC-I expression observed in *Lsd1*-null *PyMT* tumor cells (Figure 2) may therefore reduce NK cell-mediated cytotoxicity against these cells. To test this possibility, we measured NK cell cytotoxicity toward *Lsd1*-null versus *Lsd1*-wt *PyMT* tumor (organoid) cells in a co-culture system. Consistent with this hypothesis, NK cells exhibited reduced killing of *Lsd1*-null tumor cells, as indicated by decreased Calcein release ^47^ (Figure 5A-B and Figure S5).

**Figure 5.**
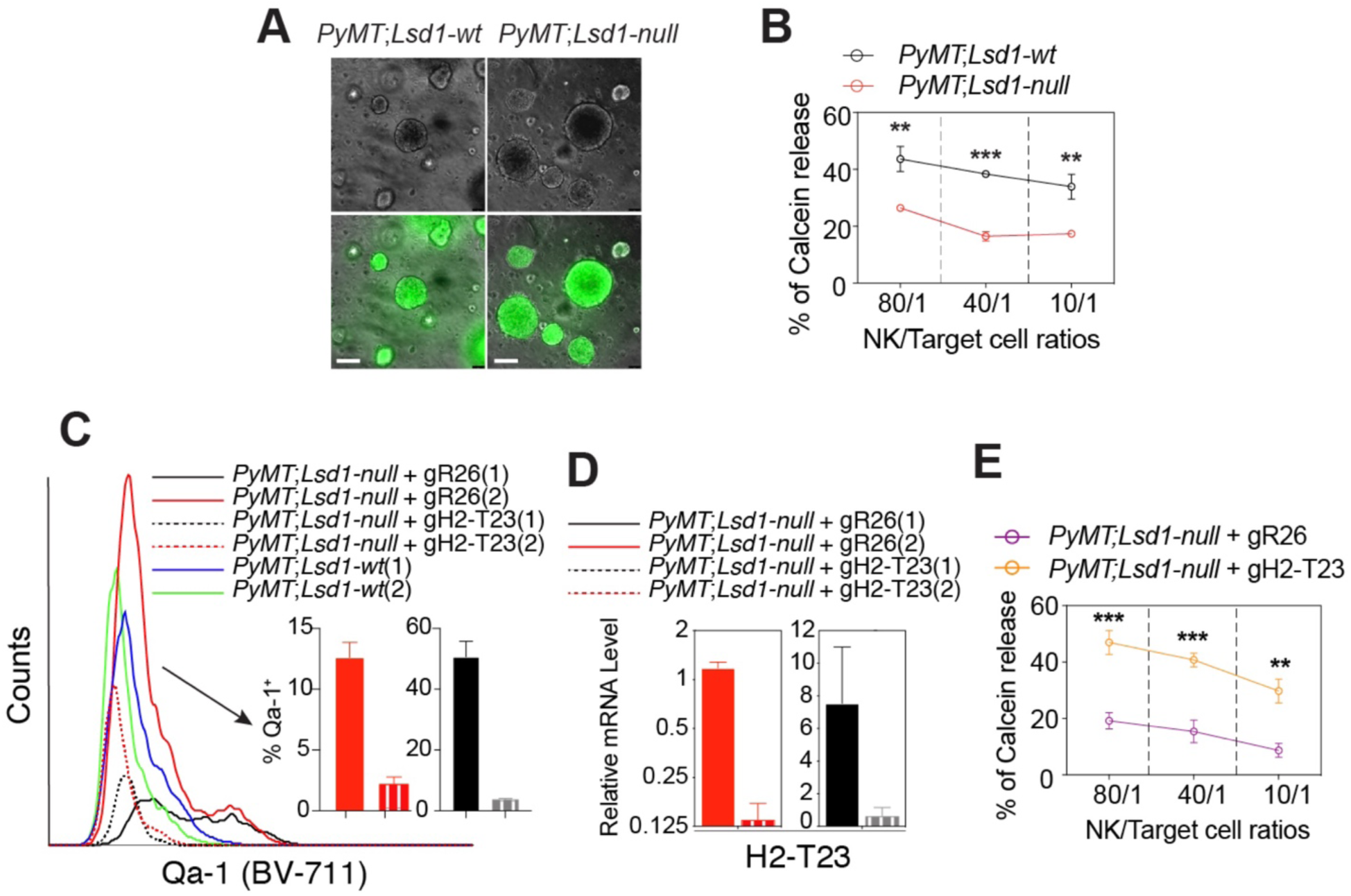
NK cells exhibited reduced cytotoxicity toward *PyMT* tumor cells with LSD1-loss. **(A**) Representative images of *Lsd1*-wt or *Lsd1*-null *PyMT* tumor cell organoids (top: phase contrast; bottom: YFP). Scale bar: 100 µm. **(B)** NK cell cytotoxicity assay based on Calcein release. **(C)** FACS analysis of Qa-1 expression from *Lsd1*-null *PyMT* tumor cell organoids with CRISPR/Cas9-based *H2-T23* knockout (gH2-T23). CRISPR/Cas9-based editing of a *Rosa26* region was used as the negative control (gR26). **(D)** qRT-PCR validation of *H2-T23* expression from *Lsd1*-null *PyMT* tumor organoid cells with *H2-T23* knockout from two independent gRNAs. **(E)** NK cell cytotoxicity assay of *Lsd1*-null *PyMT* tumor organoid cells with or without *H2-T23* knockout.

Human *HLA-E* encodes a non-classical MHC-I molecule whose functional homolog in mice is Qa-1, encoded by *H2-T23* ^48, 49^. HLA-E binds peptides derived from the conserved region of leader sequences of other MHC-I molecules ^50^, enabling its stable presentation on the cell surface. In the absence of suitable peptides, HLA-E is degraded and fails to reach the cell surface ^51^. Therefore, surface expression of HLA-E serves as an indicator of intact MHC-I expression and functional antigen processing machinery ^52^. Importantly, the inhibitory NK receptor CD94/NKG2A recognizes HLA-E/Qa-1, and their interaction suppresses NK cell activation against healthy “self” cells ^53, 54^. Consequently, stable surface expression of HLA-E/Qa-1 requires sufficient expression of both HLA-E/Qa-1 itself and other MHC-I molecules. In *Lsd1*-null *PyMT* tumor cells, both *H2-T23* and many other MHC-I genes were expressed at significantly higher levels compared with *Lsd1*-wt cells (Figure 2), suggesting that Qa-1 might be more abundantly displayed on the surface of *Lsd1*-null tumor cells and thereby inhibit NK cell activity through the CD94/NKG2A receptor. Consistent with this idea, FACS analysis confirmed elevated Qa-1 surface levels in *PyMT* tumor cells with LSD1-loss (Figure 2E-F and Figure 4C).

To determine whether increased Qa-1 expression in *Lsd1*-null *PyMT* tumor cells was responsible for the attenuated NK cell activities observed in the co-culture (Figure 5B), we disrupted Qa-1 expression in *PyMT* tumor organoid cells by CRISPR/Cas9-mediated KO of its coding gene, *H2-T23*. As a negative control, CRISPR editing targeting a neutral region within the mouse *Rosa26* locus (gR26) was used. Successful knockout of *H2-T23* was confirmed by both FACS analysis (Qa-1 protein levels) and qRT-PCR (Figure 5C-D). When *Lsd1*-null *PyMT* tumor cells lacking *H2-T23* were co-cultured with NK cells, loss of Qa-1 restored NK cell cytotoxic activity toward these tumor organoid cells (Figure 5E).

### Absence of innate immune cells abolishes the difference in metastatic potential between *Lsd1*-null and *Lsd1*-wt *PyMT* tumor cells

To determine whether NK cells contribute to the differences in metastatic potential between *Lsd1*-null and *Lsd1*-wt *PyMT* tumor cells *in vivo*, we intraductally injected these tumor cells (maintained as organoids) into the MGs of either NOD-SCID;IL2Rγ^null^ (NSG) (JAX) or SCID (CRL) recipient mice (Figure 6A). The key difference between these two recipient mouse strains is that NSG mice lack innate immune cells, including NK cells and macrophages, whereas SCID mice retain NK cells and functional macrophages ^55^.

**Figure 6.**
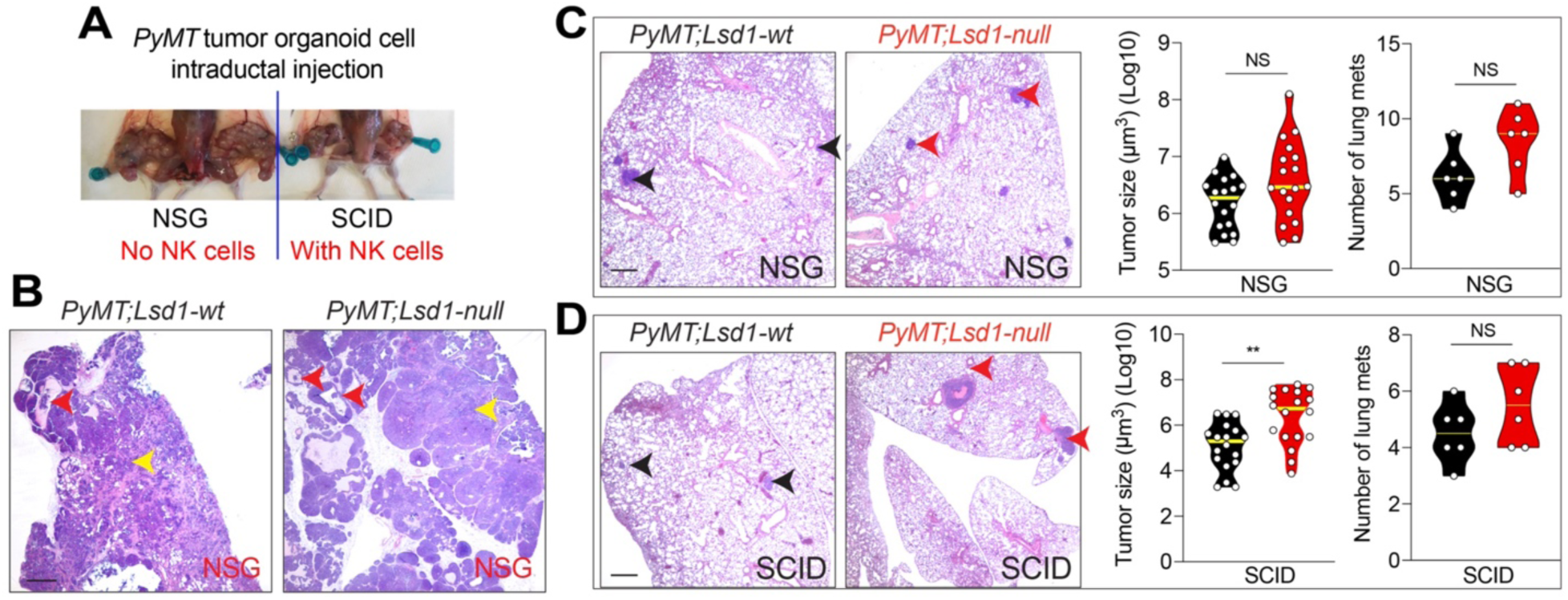
Abolishment of increased lung metastasis from *Lsd1*-null *PyMT* tumor cells in the absence of innate immune cells. **(A)** Representative images of tumors growing in MGs of NSG and SCID mice after intraductal injection of *PyMT* tumor organoid cells. **(B)** H&E pictures showing mammary tumors formed in the ducts (red arrows) and those invaded into stroma (yellow arrows), one month after intraductal injection of *Lsd1*-wt or *Lsd1*-null *PyMT* tumor cells to NSG mice. Scale bar: 500 µm. **(C)** Left: metastatic lesions formed in lungs of NSG females upon intraductal injection of *Lsd1*-wt or *Lsd1*-null *PyMT* tumor organoid cells to their MGs; middle and right: quantification of sizes and numbers of lung metastatic lesions formed in NSG recipients, respectively. Scale bar: 500 µm. **(D)** Left: metastatic lesions formed in lungs of SCID females upon intraductal injection of *Lsd1*-wt or *Lsd1*-null *PyMT* tumor organoid cells to their MGs; middle and right: quantification of sizes and numbers of lung metastatic lesions formed in SCID recipients, respectively. Scale bar: 500 µm. *P* value: *p<0.05, **p<0.01, ***p<0.005, NS = not significant, two-tailed Student’s *t*-test. Data represent mean ± SEM.

Following intraductal injection, *PyMT* tumor organoid cells formed *in situ* tumors within the mammary ducts, subsequently invading the surrounding stroma in the MGs of recipient mice (Figure 6B), and progressed to lung metastases within approximately one month (Figure 6C-D). We found that these tumor organoid cells generated similar numbers of lung metastatic lesions in both types of recipients (Figure 6C-D). However, the sizes of the metastatic lesions differed markedly between the two models. In NSG recipients, which lack NK cells, both *Lsd1*-null and *Lsd1*-wt *PyMT* tumor cells formed lung metastatic lesions of comparable size (Figure 6C). In contrast, in SCID recipients, which retain NK cells, metastatic lesions derived from *Lsd1*-wt tumor cells were significantly smaller than those derived from *Lsd1*-null tumor cells (Figure 6D).

In addition to NK cells, SCID mice also possess functional macrophages. However, macrophages have been reported to promote, rather than suppress, metastatic progression in the *PyMT* model ^56, 57^, in contrast to the anti-metastatic role of NK cells. Moreover, our analysis of *PyMT* primary tumors with induced LSD1-loss revealed a marked reduction in macrophage abundance compared with tumors retaining wt LSD1 (Figure 3B and Figure 4E). Therefore, although macrophages may still contribute to metastatic progression in the *PyMT* model, they are less likely to account for the enhanced metastasis observed in *PyMT* tumor cells with LSD1-loss or for the smaller lung metastases formed by *Lsd1*-wt tumor cells in SCID recipients. Taken together, these *in vivo* data support a model in which innate immune cells, particularly NK cells, play an important role in restricting lung metastasis of *PyMT* tumor cells and contribute to the differential metastatic capacities of *Lsd1*-null versus *Lsd1*-wt *PyMT* tumor cells.

### A negative correlation between *LSD1* and *HLE-A* expression is conserved in human luminal breast cancer cells

To extend our observation from the *PyMT* mouse model to human breast cancer, we analyzed the expression of LSD1 and MHC-I genes in human luminal breast tumors. Across human breast cancer, *LSD1* (*KDM1A*) expression showed the strongest negative correlation with *HLA-E* expression among MHC-I genes, particularly in Luminal A and HER2-enriched subtypes (Figure S6A). To further validate this relationship, we analyzed the largest publicly available breast cancer cohort, METABRIC ^58, 59^. Among 1,445 estrogen receptor-positive (ER^+^) luminal breast tumors in this cohort with microarray data, the top 165 and 143 cases with the highest and lowest expression levels of *LSD1* (*KDM1A*) were classified as *LSD1*-high and *LSD1*-low groups, respectively. We found that tumors in the *LSD1*-low group exhibited significantly higher expression levels of multiple MHC-I genes, most notably the non-classical MHC-I gene *HLA-E* (Figure 7A). Importantly, GSEA of the expression profiling data revealed that many of the top enriched gene sets in *LSD1*-low tumors (relative to *LSD1*-high tumors) were immune-related, including pathways associated with IFN signaling (Figure 7B and Figure S6B). Lastly, in the human ER^+^ luminal breast cancer cell line MCF7 ^18^, knockdown of *LSD1* led to upregulation of several MHC-I genes, with *HLA-E* showing the most prominent increase (Figure S6C).

**Figure 7.**
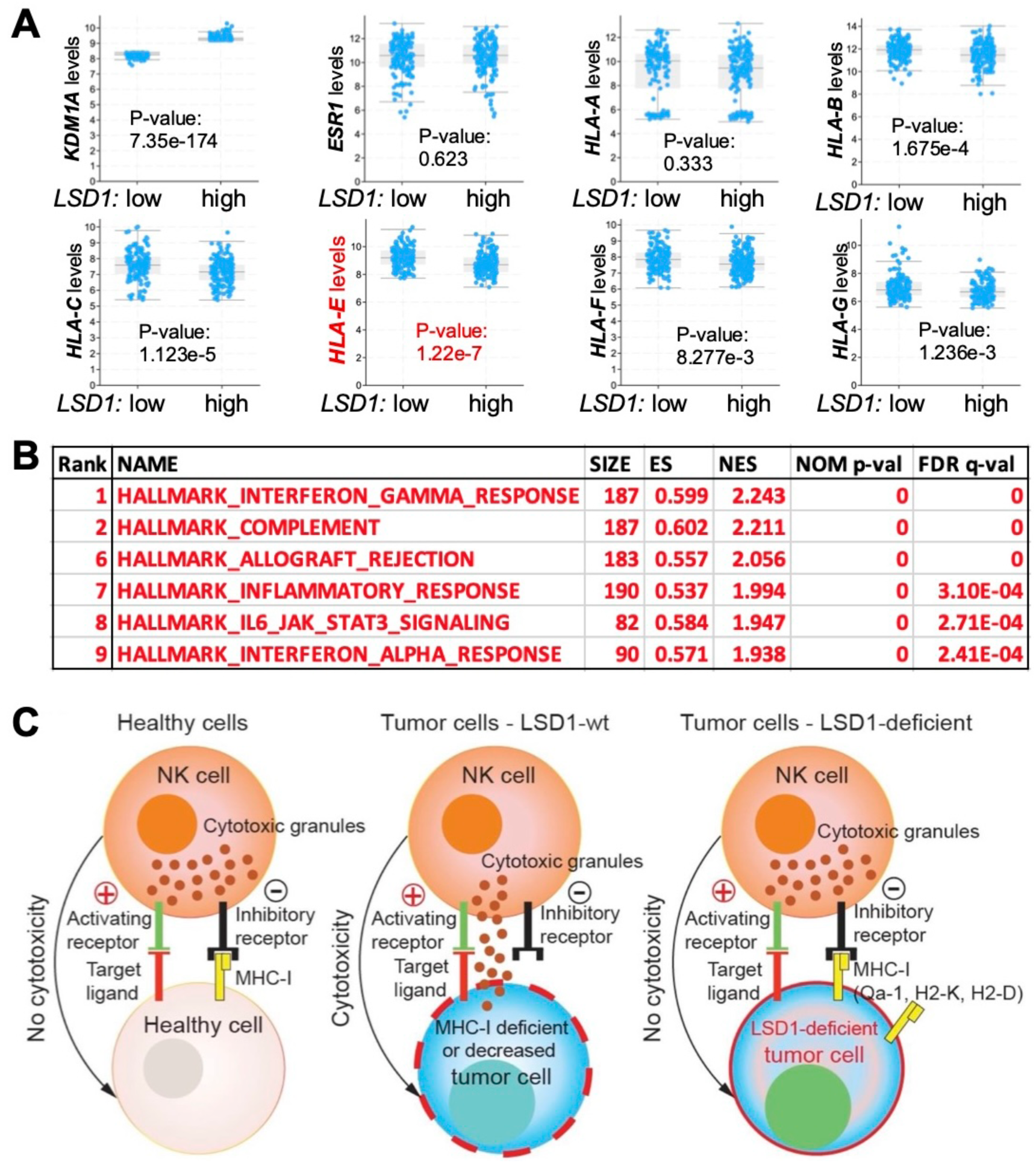
Negative correlation of *LSD1* with *HLA-E* in human luminal breast cancer. **(A)** Top 165 and 143 cases with the highest and lowest *LSD1* (*KDM1A*) expression levels among 1,445 ER^+^ breast tumors with microarray data in the METABRIC cohort were assigned to *LSD1*-high and *LSD1*-low groups, respectively. These two groups have similar *ESR1* expression levels, but the *LSD1*-low group has significantly higher expression levels of most MHC-I genes, particularly *HLA-E*; the analysis was based on cbioportal. **(B)** GSEA showing top enriched cancer hallmark-gene sets (from the GSEA MSigDB collection, only immune-related gene sets are shown) in *LSD1*-low group (in relation to *LSD1*-high group). ES: enrichment score; NES: normalized enrichment score; NOM p-val: nominal p-value; FDR q-val: false discovery rate q-value. See also Figure S6B. **(C)** Proposed model for LSD1 ablation and attenuated NK cell-mediated anti-tumor immunity. Schematic diagram summarizing the attenuated cytotoxicity of NK cells toward LSD1-deficient tumor cells with increased expression of MHC-I molecules.

Together, our *in vivo* and *in vitro* data support a model in which LSD1 ablation in *PyMT* tumor cells increases their immunogenicity and elevates MHC-I expression, thereby reducing NK cell-mediated anti-tumor immunity (Figure 7C). The resulting attenuation of NK cell-mediated tumor surveillance may contribute to the enhanced lung metastatic growth observed in *Lsd1*-null *PyMT* tumors. Notably, the similar upregulation of MHC-I genes, particularly *HLA-E*, in human luminal breast cancer cells with reduced LSD1 expression (or LSD1 deficiency) suggests that a comparable mechanism may operate in human disease, potentially enabling tumor cells to evade NK cell-mediated elimination and thereby promoting metastasis.

## Discussion

In this study, we demonstrate that, consistent with previous reports ^30, 60^, genetic ablation of LSD1 increases tumor cell immunogenicity. However, when the immune system fails to effectively eliminate these tumor cells, the resulting prolonged immune activation can lead to chronic inflammation and reprogramming of the tumor immune microenvironment. One consequence of this altered immune landscape is increased expression of MHC-I genes in tumor cells lacking LSD1, which attenuates NK cell-mediated anti-tumor immunity (Figure 7C). As NK cells play an essential role in restricting metastatic progression (Figure 3F) ^43^, these changes in MHC-I expression as well as NK cell abundance, maturation, and activity may contribute to the increased lung metastatic burden observed in *PyMT* mice following LSD1-loss.

This conclusion is supported by our *in vivo* transplantation experiment, in which differences in lung metastasis between *Lsd1*-null and *Lsd1*-wt *PyMT* tumor cells were observed in SCID recipients (which retain NK cells) but not in NSG recipients (which lack NK cells). In SCID mice, we observed a significant difference in the size of lung metastatic lesions (Figure 6D), although not in their number. The absence of differences in lesion number may reflect the relatively short duration of the experiment (∼1 month), as the rapid growth of primary tumors in the injected MGs required early euthanasia of recipient mice. Of note, the involvement of NK cells in this phenotype does not exclude contributions from other immune cell populations to the increased lung metastasis associated with LSD1 ablation. Previous studies have shown that macrophages play a key role in *PyMT* tumor metastasis ^57^, and that CD4^+^ T cells promote lung metastasis of *PyMT* tumors by enhancing the pro-tumorigenic properties of macrophages ^56^. Although we observed an increased CD4⁺ T cell population in *PyMT* tumors with LSD1-loss, the macrophage population was profoundly reduced (Figure 3B and Figure 4E), suggesting that CD4⁺ T cells or macrophages are unlikely to be the primary immune cell populations responsible for the enhanced metastasis observed following LSD1 ablation.

In a previous study using the 4T1 mammary tumor syngeneic transplantation model, cotreatment with anti-PD-1 antibody and the LSD1 inhibitor HCI-2509 was reported to reduce tumor burden. However, closer examination of the data indicates that treatment with HCI-2509 alone resulted in a modest increase in lung metastasis (∼30% increase in lesion size) ^60^. In a preliminary study, we similarly treated *PyMT* female mice with HCI-2509 for ∼2 months (beginning at ∼4 weeks of age, comparable to the timing of induced *Lsd1* deletion in our genetic models) and observed significant increases in both the numbers and sizes of lung metastases. In these inhibitor-treated mice, analysis of *PyMT* tumors in the MGs also revealed increased numbers of Qa-1^+^, H2-K^+^ and H2-D^+^ tumor cells, accompanied by a reduction in the overall NK cell population and an increase in the immature NK cell subset among CD45⁺ leukocytes. Together, these observations suggest that pharmacological inhibition of LSD1, particularly when administered over a prolonged period, may also promote mammary tumor metastasis, at least in part through attenuation of NK cell-mediated anti-tumor immunity.

One important caveat is that, whereas our genetic approaches disrupt LSD1 specifically in *PyMT* tumor cells, pharmacological inhibition of LSD1 is expected to affect multiple cell types, including immune cells such as NK cells. Of note, recent work has shown that different classes of LSD1 inhibitors exert distinct effects on NK cells: catalytic inhibitors that block the demethylase activity of LSD1 appear to have minimal impact on NK cell function, whereas scaffolding inhibitors that disrupt LSD1-associated epigenetic complexes can impair NK cell cytolytic activity ^61^. HCI-2509 (also known as SP-2509) is a scaffolding inhibitor of LSD1, and thus the increased metastasis observed following HCI-2509 treatment could partly reflect its direct inhibitory effects on NK cell function. Nevertheless, these findings further support a role for impaired NK cell-mediated anti-tumor immunity in the enhanced metastasis associated with LSD1-loss. This impairment may arise through increased HLA-E(Qa-1)-CD94/NKG2A inhibitory signaling in tumor cells, direct inhibition of NK cell function by certain classes of LSD1 inhibitors, or a combination of both mechanisms. Future studies comparing different classes of LSD1 inhibitors *in vivo* will be important for disentangling these NK cell-related mechanisms in breast cancer metastasis.

Compared with previous studies ^30, 60^, which relied on transplantation models typically lasting only one to two months, our work employs an autochthonous breast cancer model in which tumors evolve under physiological conditions for more than two months. This system therefore captures the longer-term consequences of LSD1 disruption during tumor progression. Our findings suggest that although short-term LSD1 inhibition may enhance tumor cell immunogenicity and thereby improve the efficacy of ICB-based therapies, prolonged disruption of LSD1 may instead promote chronic inflammation and potentially increase metastatic risk. We identify reduced NK cell-mediated anti-tumor immunity as a key mechanism contributing to this increased metastatic potential, likely mediated in part through activation of the HLA-E(Qa-1)-CD94/NKG2A inhibitory pathway.

The recent development of therapeutic antibodies targeting NKG2A, which can enhance anti-tumor immunity by unleashing both NK and T cell activity ^40^, provides an intriguing potential strategy. We propose that combining LSD1 inhibitors, particularly catalytic inhibitors, with anti-NKG2A antibody therapy may both enhance checkpoint inhibitor-based immunotherapy and restore NK cell-mediated tumor surveillance, including NK cell-dependent suppression of metastasis.

## Materials and methods

### Mouse models

The *Lsd1^L/L^* (B6.129-*Kdm1a^tm^*^1^*^.1Sho^*/J) conditional knockout mouse line was obtained from Dr. Stuart Orkin (Harvard Medical School). *Rosa26-LSL-YFP* (*R26Y*) [*B6.129X1-Gt(ROSA)26Sor^tm^*^1^(EYFP)*^Cos^*/J, Strain #: 006148] reporter mice, *MMTV-PyMT* [*FVB/N-Tg(MMTV-PyMT)634Mul/J*, Strain #: 002374, or *B6.FVB-Tg(MMTV-PyVT)634Mul/LellJ*, Strain #: 022974] and *K8-CreER* [*STOCK Tg(Krt8-cre/ERT2)17Blpn/J*, Strain #: 017947] transgenic mice, *Cd8a^-/-^*[*B6.129S2-Cd8atm1Mak/J*, Strain #: 002665] and *Il2rg^-/-^* [*B6.129S4-Il2rgtm1Wjl/J*, Strain #: 003174] mice were obtained from The Jackson Laboratory (JAX) (Bar Harbor, ME). For this study, we first bred *MMTV-PyMT* mice (FVB/N background) with *Lsd1^L/L^;R26Y* mice (B6.129 background) and generated a heterozygous strain (*MMTV-PyMT;Lsd1^L/+^;R26Y* or *Lsd1^L/+^;R26Y* littermates). The heterozygous *MMTV-PyMT;Lsd1^L/+^;R26Y* males and *Lsd1^L/+^;R26Y* females (mixed FVB/B6.129 background) were set up as breeding pairs to generate the *MMTV-PyMT;Lsd1^L/L^;R26Y* experimental and *MMTV-PyMT;R26Y* control mice for *Ad-K8-Cre* intraductal injection. For the *K8-CreER*-based model, we first crossed *K8-CreER* mice (B6 background) to *MMTV-PyMT;Lsd1^L/L^;R26Y* mice (FVB/B6.129 mixed) to generate *MMTV-PyMT;K8-CreER;Lsd1^L/+^;R26Y* and *K8-CreER;Lsd1^L/+^;R26Y* mice, which were then intercrossed to generate *MMTV-PyMT;K8-CreER;Lsd1^L/L^;R26Y* experimental and *MMTV-PyMT;K8-CreER;R26Y* control mice. Immunodeficient NOD-SCID;IL2Rγ^null^ (NSG, JAX Strain #: 005557) mice and SCID mice were obtained from JAX and Charles River Laboratories (CRL) (Wilmington, MA), respectively. To target luminal MECs in *MMTV-PyMT;Lsd1^L/L^;R26Y* or *MMTV-PyMT;R26Y* female mice, mice were anesthetized and *Ad-K8-Cre* adenovirus [diluted in injection medium (DMEM supplemented with 0.1% Bromophenol blue and 0.01M CaCl_2_)] were introduced into mammary ducts of their #4 inguinal glands via intraductal injection ^32, 62^. For *MMTV-PyMT;K8-CreER*-based females, to induce CreER activity, tamoxifen (2 mg per mouse, one injection) was introduced into adult mice (8 weeks of age) by intraperitoneal injection ^63^. The tumor growth was evaluated at every other day; lungs of experimental and control mice were dissected at the experimental end point and fixed in paraffin, followed by histological processing and hematoxylin and eosin (H&E) staining at the Dana-Farber/Harvard Cancer Center (DF/HCC) Rodent Histopathology Core. The whole lung tissue was sectioned sagittally and at least two sections per mouse were evaluated; each experiment (including 1-3 experimental and control mice) was repeated at least eight times (total mice: *Ad-K8-Cre*: experimental n=20, control n=18; *K8-CreER*: experimental n=15, control n=10) to evaluate the statistical significance between the experimental and control groups. The blinded evaluation of metastasis was conducted by a pathologist (R.T. Bronson). All animal experiments were approved by the Institutional Animal Care and Use Committee (IACUC) of Brigham and Women’s Hospital (2016N000363, 2020N000122).

### Mammary tumor cell preparation

Mammary tumors were dissected and minced, and then incubated in digestion medium (DMEM/F12 with 2% Penicillin/Streptomycin, 0.1 mg ml^-1^ Gentamicin, 0.6% Nystatin, 2 mg ml^-1^ Collagenase A, 0.096 mg ml^-1^ Hyaluronidase) at 37°C with shaking for 1-1.5 hours. After digestion, the cells/tissues were treated sequentially with 0.25% trypsin/EDTA (37°C, 2 minutes), 5 mg ml^-1^ dispase with DNaseI (0.1mg ml^-1^, Sigma, St. Louis, MO) (37°C, 5 minutes), cold red blood cell (RBC) lysis buffer (on ice, 2-3 minutes). Between each treatment step, cells/tissues were washed with 1x PBS with 5% FCS. After treatment with the RBC lysis buffer, cells/tissues were filtered through a 40 μm cell strainer and washed with 1x PBS/5% FCS, to obtain single cell suspension ^64^.

### Flow cytometry

Single cell suspensions obtained from *MMTV-PyMT* tumors with or without LSD1-loss were analysed by flow cytometric (FACS) analysis, evaluating the immune cell populations. The FACS analysis and cell sorting were performed with the BD FACSymphony and the BD FACSAria sorter (BD Biosciences), respectively. Antibodies targeting immune cells used in FACS analysis were purchased from eBiosciences: CD11b-PE-Cyanine7 (clone M1/70, Cat. No. 25-0112-82); CD19-PE (clone eBio1D3 (1D3), Cat. No. 12-0193-81); H2-D-BV786 (clone KH117, Cat. No. 744665), or from BD Biosciences: Qa-1(b)-BV711 (clone 6A8.6F10.1A6, Cat. No. 744389), or from BioLegend: CD45-PerCP/Cyanine5.5 (Clone 30-F11, Cat. No. 103131), Ly-6G-APC/Cy7 (Clone 1A8, Cat. No. 127623), F4/80-Brilliant Violet 605™ (Clone BM8, Cat. No. 123133), NK-1.1-Brilliant Violet 650™ (Clone PK136, Cat. No. 108735), CD3-APC (Clone 17A2, Cat. No. 100235), CD4-FITC (Clone GK1.5, Cat. No. 100405); CD8-APC (Clone 17A2, Cat. No. 100235); CD107a-Alexa Fluor® 700 (Clone 1D4B, Cat. No. 121627); H2-K-PE (Clone AF6-88.5, Cat. No. 116507); CD69 (APC-Cyanine7, clone H1.2F3, Cat. No. 104525), and NKp46 (PE, clone 29A1.4, Cat. No. 137603). Data analyses were performed using FlowJo 10.6.0 (FlowJo LLC ©).

### Mammary organoid culture and intraductal injection

The organoid culture was established by seeding YFP^+^ cells sorted from MGs of *MMTV-PyMT;Lsd1^L/L^;R26Y* or *MMTV-PyMT;R26Y* female mice ∼2 weeks after intraductal injection of *Ad-K8-Cre* and was conducted by following a protocol we reported previously ^65^. For transplantation, *MMTV-PyMT* tumor organoid cells (mixed background of C57BL/6 and FVB) with or without LSD-loss (20,000 cells) were injected into each #4 inguinal glands of NSG (JAX) or SCID (CRL) mice at ∼5 weeks of age via intraductal injection (n=6) ^32, 62^. *MMTV-PyMT* tumor organoid cells (pure C57BL/6 background) (20,000 cells) were injected into each #4 inguinal glands of syngeneic *Cd8a^-/-^*(JAX, strain# 002665) (n=5) or *Il2rg^-/-^* (JAX, strain# 003174) (n=4) or wildtype (JAX, strain #: 000664) (n=5) female mice at ∼8 weeks of age via intraductal injection. At the end point following tumor size evaluation, mice were euthanized and lung tissues from each mouse were fixed, embedded in paraffin; the resulting paraffin sections were processed for H&E-staining at the DF/HCC Rodent Histopathology Core. The blinded evaluation of metastasis was conducted by a pathologist (R.T. Bronson).

### NK cell isolation and Calcein-AM cytotoxicity assay

The general NK cell cytotoxicity assay was completed as described ^66^. Specifically, splenic NK cells from naïve wt mice were isolated according to the manufacturer’s instruction (NK Cell Isolation Kit, mouse; #130-115-818; Miltenyi Biotec) and suspended in complete NK-cell buffer (phenol-red free RPMI 1640 supplemented with 10% FBS, β-mercaptoethanol, non-essential amino acids, L-glutamine, sodium pyruvate, all from Gibco). The purity of the enriched NK cells was further assessed by FACS using antibodies CD45 (PerCP/Cyanine5.5, clone 30-F11, 103131, BioLegend), CD11b (PE-Cyanine7, clone M1/70, 25-0112-82, eBiosciences), CD3 (APC, clone 17A2, 100235, BioLegend), CD69 (APC-Cyanine7, clone H1.2F3, 104525, BioLegend), and NKp46 (PE, clone 29A1.4, 137603, BioLegend). *MMTV-PyMT* tumor organoid cells with or without LSD-loss were labeled with 10-15 μM Calcein-AM (5016952; FISHER SCIENTIFIC LLC) for 20-30 minutes at 37°C/5% CO2, allowing Calcein-AM to enter organoid cells. The purified NK cells and the target organoid cells were mixed at various ratios (80:1, 40:1, 10:1) in at least three replicates in a round-bottom, 96-well plate, and incubated at 37°C/5% CO2 for 4 hours. The spontaneous (i.e., only target organoid cells in culture medium) and maximum release (only target organoid cells in medium plus 2% Triton X-100 buffer) groups were also included. Calcein release was quantified by harvesting 100 μL of cell-free supernatant to a new plate and measuring fluorescent emission at the appropriate wavelength (excitation filter: 485±9 nm; band-pass filter: 530±9 nm) using a multilabel counter (VICTOR3, Perkin Elmer, San Diego, CA, USA). Cytotoxicity data were expressed as percent release relative to the spontaneous (target cells alone) and maximum release (2% Triton X-100 treated cells) for a particular target with the equation: % of Calcein release = (Test release-spontaneous release)/ (Maximum release-spontaneous release) * 100%.

### Immunofluorescence (IF) staining

IF staining was performed on sections from mammary tumors that were fixed in 10% formalin (Fisher Scientific, Hampton, NH) and embedded in paraffin. The IF staining was performed as described ^33^. Antigen retrieval (Citrate buffer pH 6.0, 20 min boil in microwave oven) was performed before blocking. Primary antibodies used included: anti-GFP (ab290, 1:500 or ab6673, 1:200, Abcam, Cambridge, UK), anti-Keratin 8 (K8) (MMS-162P, 1:200, Covance), The secondary antibodies used were goat anti-mouse IgG conjugated with AF647 (A31571, 1:250), goat anti-rabbit IgG conjugated with AF488 (A11008, 1:250) or with AF594 (A11037, 1:250). Slides were counterstained with DAPI (1 μg ml^-1^).

### RNA isolation and bulk RNA sequencing

Single-cell suspensions prepared from mammary tumors post *Ad-K8-Cre* injection were obtained as described in the “Mammary tumor cell preparation” procedure. Cell preps were stained with DAPI and DAPI^-^YFP^+^ cells were sorted in a FACS Aria II (BD Biosciences). Total RNA was extracted using the RNeasy Micro Kit (QIAGEN, Cat# 74004). Libraries were prepared by the Dana-Farber Cancer Institute (DFCI) Molecular Biology Cores, using the Low Input mRNA Library (Clontech SMARTer) v4, following the vendor’s protocol. Library quality was checked using High Sensitivity D1000 reagents in an Agilent Tape Station. Libraries were pooled and sequenced in an Illumina NextSeq500 (Single-End 75bp reads per sample). A Visualization Pipeline for RNAseq (Viper) analysis tool, developed by the Center for Functional Cancer Epigenetics (CFCE) at DFCI, was used to generate standard outputs.

### Quantitative RT-PCR (qRT-PCR)

Organoids were dissociated with TrypLE and pelleted by centrifugation for 5 minutes at 1,000 rpm, followed by RNA extraction with either TRIzol (Thermo Fisher Scientific, 15596026) or the Allprep DNA/RNA mini/micro kits (Qiagen) according to the manufacturer’s protocols. cDNA was generated with an iScript cDNA synthesis kit (Bio-RAD, 170-8891). Quantitative RT-PCR was performed using FastStart SYBR Green Master (Roche, 04913850001, IN).

The primer sequences are listed below for qRT-PCR ^37, 38, 67^:

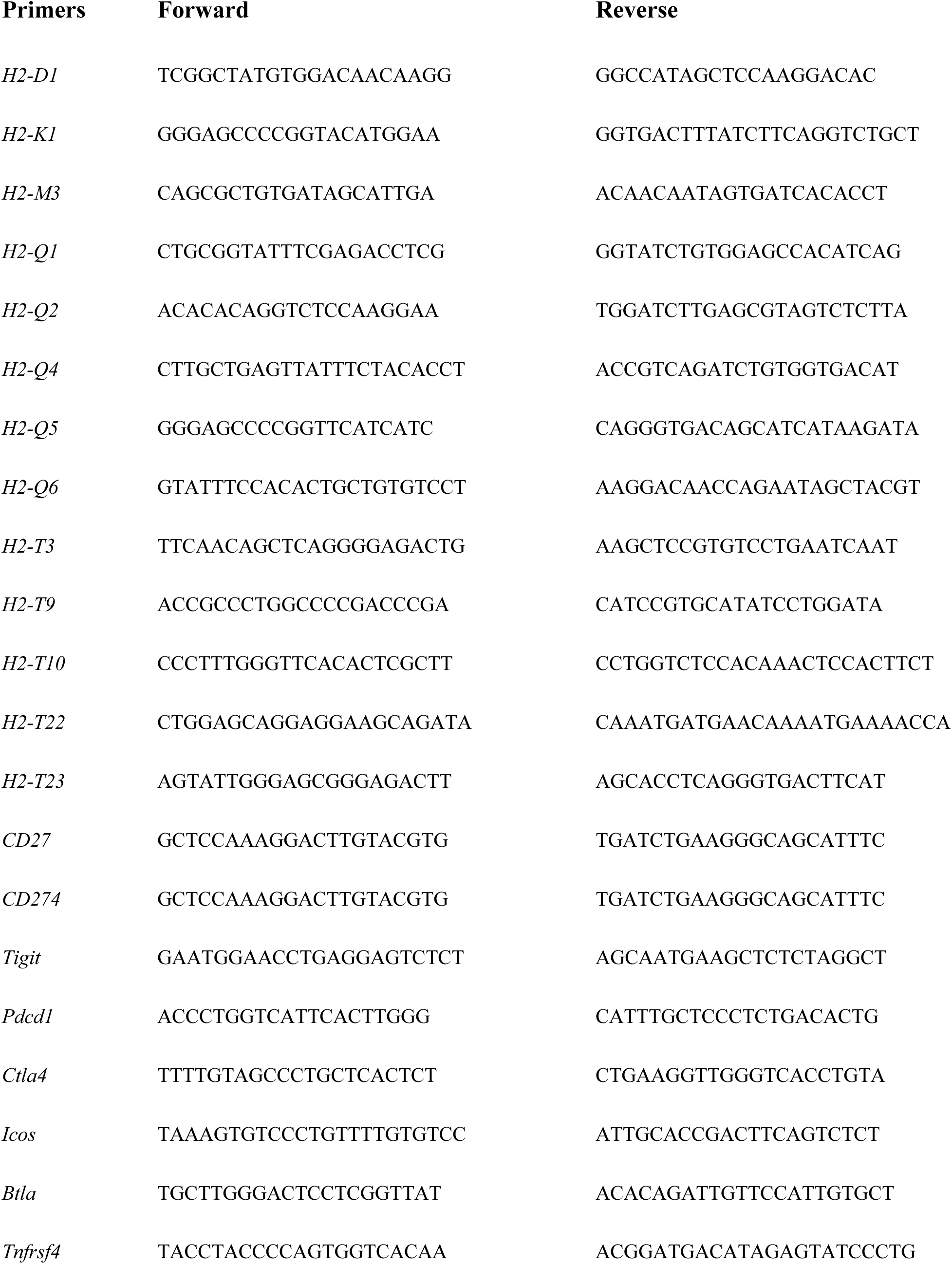
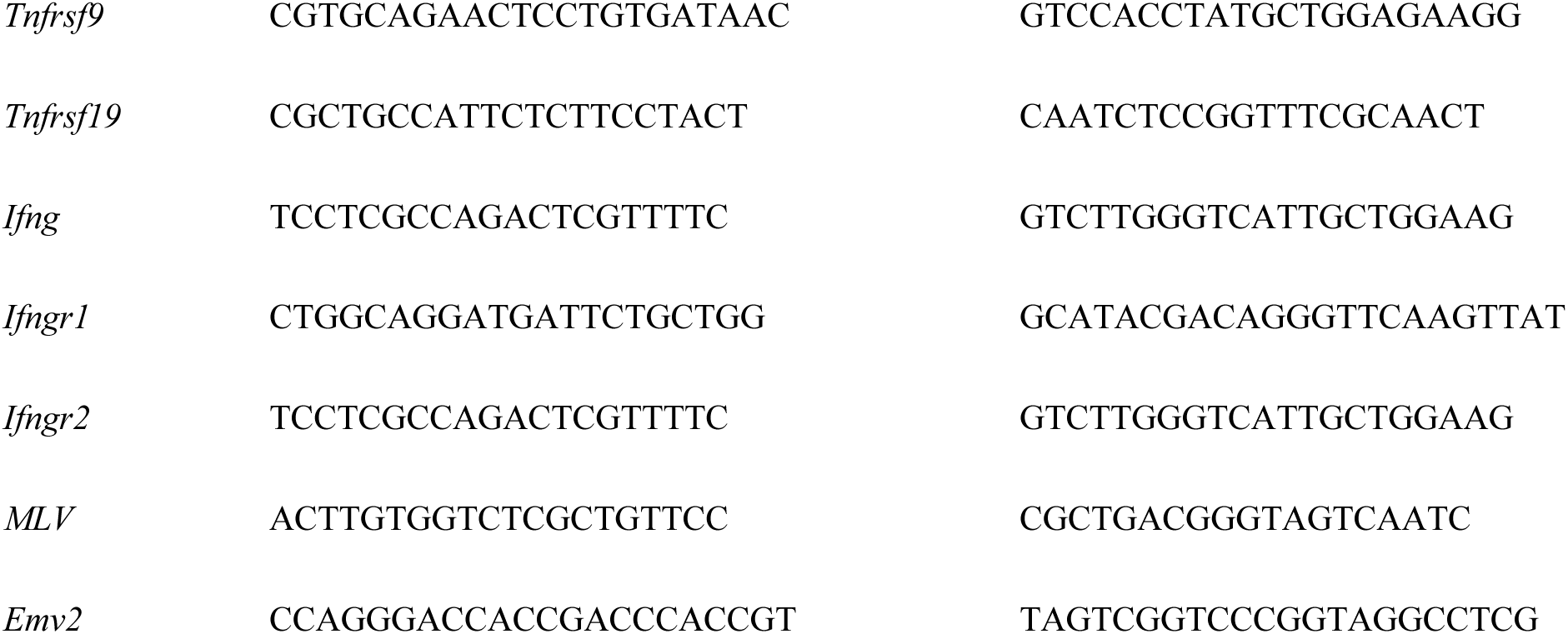

### CRISPR/Cas9 lentiviral vectors and organoid manipulation

The lentiCRISPR-v2 plasmid was from Addgene (No. 52961, a gift from Dr. Feng Zhang). Two individual guide RNAs (gRNAs) targeting *H2-T23* or *Kdm1a* were designed based on http://crispr.mit.edu (with a high-quality score of above 90) (g*H2-T23*-1: CTTGTGCTTAGAGATCTGTG; g*H2-T23*-2: TACTACAATCAGAGTAACGA; g*Kdm1a*-1: TGTAAGGCGCTTCCAGCTGC; g*Kdm1a*-1: TCAACTTCGGCATCTACAAG). LentiGuide vectors constructed with gRNAs targeting the mouse *Rosa26* region of mouse (gR26) was applied as the negative control ^65^. Cloning of gRNAs in lentiCRISPR-v2 plasmid was performed as described ^68^. All vectors were validated by Sanger sequencing. We produced concentrated lentiviral stocks, pseudo-typed with the VSV-G envelope protein, by transient transfection of their corresponding plasmids into 293T cells as previously described ^69^.

The procedure for organoid transfection was developed with modifications from the protocol described previously ^70^. Specifically, following one day of culture in full medium, as described above, tumor cell organoids were infected with lentivirus for 2-3 days before selection was initiated in full culture medium containing puromycin (1 µg ml^-1^) for ∼one week. Stably infected and edited organoid cells were expanded in full medium. The NK cell cytotoxic assay toward transfected tumor organoid cells was conducted as described in the section of “NK cell isolation and Calcein-AM cytotoxicity assay”.

### Statistics

Student’s *t*-test (two-tailed) was used to calculate the *P* values. Data were reported as mean ± SEM. No statistical method was used to pre-determine the sample size for mice. No randomization or blinding was used in the *in vivo* studies.

### Data and materials availability

The RNAseq data has been deposited in the NCBI GEO database (http://www.ncbi.nlm.nih.gov/geo/) under accession code GSE140820.

## Supplemental material

This article includes six supplementary figures (Figures. S1-S6).

## Author contributions

D.X. and S.H. conducted experiments, analyzed data, and wrote the manuscript. A.H. conducted experiments and analyzed data. G.Q. conducted experiments. R.T.B. performed histology evaluation. Z.L. designed experiments, analyzed data, and wrote the manuscript.

## Acknowledgments

We are grateful to Yiling Qiu, Yidong Lin and Chad Araneo for their expert technical assistance with FACS. We thank Dr. Stuart Orkin for the *Lsd1^L/L^* conditional knockout mouse line. This research was supported by a Breakthrough Award from U.S. Department of Defense (W81XWH-15-1-0100) and by NIH/NCI R01 grants (R01CA222560, R01CA248306) to ZL. AH is supported by St. Baldrick’s Foundation (585350). The authors declare no competing financial interests.

## References

1 Chatterjee A, Rodger EJ, Eccles MR. Epigenetic drivers of tumourigenesis and cancer metastasis. Semin Cancer Biol 2018; 51: 149–159.

2 Patel SA, Vanharanta S. Epigenetic determinants of metastasis. Mol Oncol 2017; 11: 79–96.

3 Vogelstein B, Papadopoulos N, Velculescu VE, Zhou S, Diaz LA, Jr., Kinzler KW. Cancer genome landscapes. Science 2013; 339: 1546–1558.

4 Turajlic S, Swanton C. Metastasis as an evolutionary process. Science 2016; 352: 169–175.

5 Yates LR, Knappskog S, Wedge D, Farmery JHR, Gonzalez S, Martincorena I, Alexandrov LB, Van Loo P, Haugland HK, Lilleng PK, Gundem G, Gerstung M, Pappaemmanuil E, Gazinska P, Bhosle SG, Jones D, Raine K, Mudie L, Latimer C, Sawyer E, Desmedt C, Sotiriou C, Stratton MR, Sieuwerts AM, Lynch AG, Martens JW, Richardson AL, Tutt A, Lonning PE, Campbell PJ. Genomic Evolution of Breast Cancer Metastasis and Relapse. Cancer Cell 2017; 32: 169–184 e167.

6 Bertucci F, Ng CKY, Patsouris A, Droin N, Piscuoglio S, Carbuccia N, Soria JC, Dien AT, Adnani Y, Kamal M, Garnier S, Meurice G, Jimenez M, Dogan S, Verret B, Chaffanet M, Bachelot T, Campone M, Lefeuvre C, Bonnefoi H, Dalenc F, Jacquet A, De Filippo MR, Babbar N, Birnbaum D, Filleron T, Le Tourneau C, Andre F. Genomic characterization of metastatic breast cancers. Nature 2019; 569: 560–564.

7 Priestley P, Baber J, Lolkema MP, Steeghs N, de Bruijn E, Shale C, Duyvesteyn K, Haidari S, van Hoeck A, Onstenk W, Roepman P, Voda M, Bloemendal HJ, Tjan-Heijnen VCG, van Herpen CML, Labots M, Witteveen PO, Smit EF, Sleijfer S, Voest EE, Cuppen E. Pan-cancer whole-genome analyses of metastatic solid tumours. Nature 2019; 575: 210–216.

8 Canadas I, Thummalapalli R, Kim JW, Kitajima S, Jenkins RW, Christensen CL, Campisi M, Kuang Y, Zhang Y, Gjini E, Zhang G, Tian T, Sen DR, Miao D, Imamura Y, Thai T, Piel B, Terai H, Aref AR, Hagan T, Koyama S, Watanabe M, Baba H, Adeni AE, Lydon CA, Tamayo P, Wei Z, Herlyn M, Barbie TU, Uppaluri R, Sholl LM, Sicinska E, Sands J, Rodig S, Wong KK, Paweletz CP, Watanabe H, Barbie DA. Tumor innate immunity primed by specific interferon-stimulated endogenous retroviruses. Nat Med 2018; 24: 1143–1150.

9 Hurst TP, Magiorkinis G. Activation of the innate immune response by endogenous retroviruses. J Gen Virol 2015; 96: 1207–1218.

10 Grandi N, Tramontano E. Human Endogenous Retroviruses Are Ancient Acquired Elements Still Shaping Innate Immune Responses. Front Immunol 2018; 9: 2039.

11 Shi Y, Lan F, Matson C, Mulligan P, Whetstine JR, Cole PA, Casero RA. Histone demethylation mediated by the nuclear amine oxidase homolog LSD1. Cell 2004; 119: 941–953.

12 Lee MG, Wynder C, Cooch N, Shiekhattar R. An essential role for CoREST in nucleosomal histone 3 lysine 4 demethylation. Nature 2005; 437: 432–435.

13 Wang Y, Zhang H, Chen Y, Sun Y, Yang F, Yu W, Liang J, Sun L, Yang X, Shi L, Li R, Li Y, Zhang Y, Li Q, Yi X, Shang Y. LSD1 is a subunit of the NuRD complex and targets the metastasis programs in breast cancer. Cell 2009; 138: 660–672.

14 Metzger E, Wissmann M, Yin N, Muller JM, Schneider R, Peters AH, Gunther T, Buettner R, Schule R. LSD1 demethylates repressive histone marks to promote androgen-receptor-dependent transcription. Nature 2005; 437: 436–439.

15 Wissmann M, Yin N, Muller JM, Greschik H, Fodor BD, Jenuwein T, Vogler C, Schneider R, Gunther T, Buettner R, Metzger E, Schule R. Cooperative demethylation by JMJD2C and LSD1 promotes androgen receptor-dependent gene expression. Nat Cell Biol 2007; 9: 347–353.

16 Garcia-Bassets I, Kwon YS, Telese F, Prefontaine GG, Hutt KR, Cheng CS, Ju BG, Ohgi KA, Wang J, Escoubet-Lozach L, Rose DW, Glass CK, Fu XD, Rosenfeld MG. Histone methylation-dependent mechanisms impose ligand dependency for gene activation by nuclear receptors. Cell 2007; 128: 505–518.

17 Choi HJ, Park JH, Park M, Won HY, Joo HS, Lee CH, Lee JY, Kong G. UTX inhibits EMT-induced breast CSC properties by epigenetic repression of EMT genes in cooperation with LSD1 and HDAC1. EMBO reports 2015; 16: 1288–1298.

18 Hu X, Xiang D, Xie Y, Tao L, Zhang Y, Jin Y, Pinello L, Wan Y, Yuan GC, Li Z. LSD1 suppresses invasion, migration and metastasis of luminal breast cancer cells via activation of GATA3 and repression of TRIM37 expression. Oncogene 2019; 38: 7017–7034.

19 Lim S, Janzer A, Becker A, Zimmer A, Schule R, Buettner R, Kirfel J. Lysine-specific demethylase 1 (LSD1) is highly expressed in ER-negative breast cancers and a biomarker predicting aggressive biology. Carcinogenesis 2010; 31: 512–520.

20 Serce N, Gnatzy A, Steiner S, Lorenzen H, Kirfel J, Buettner R. Elevated expression of LSD1 (Lysine-specific demethylase 1) during tumour progression from pre-invasive to invasive ductal carcinoma of the breast. BMC Clin Pathol 2012; 12: 13.

21 Huang Y, Vasilatos SN, Boric L, Shaw PG, Davidson NE. Inhibitors of histone demethylation and histone deacetylation cooperate in regulating gene expression and inhibiting growth in human breast cancer cells. Breast Cancer Res Treat 2012; 131: 777–789.

22 Wu Y, Wang Y, Yang XH, Kang T, Zhao Y, Wang C, Evers BM, Zhou BP. The deubiquitinase USP28 stabilizes LSD1 and confers stem-cell-like traits to breast cancer cells. Cell Rep 2013; 5: 224–236.

23 Zhang X, Lu F, Wang J, Yin F, Xu Z, Qi D, Wu X, Cao Y, Liang W, Liu Y, Sun H, Ye T, Zhang H. Pluripotent stem cell protein Sox2 confers sensitivity to LSD1 inhibition in cancer cells. Cell Rep 2013; 5: 445–457.

24 Kitamura T, Qian BZ, Pollard JW. Immune cell promotion of metastasis. Nat Rev Immunol 2015; 15: 73–86.

25 Emens LA. Breast Cancer Immunotherapy: Facts and Hopes. Clin Cancer Res 2018; 24: 511–520.

26 Sharma P, Hu-Lieskovan S, Wargo JA, Ribas A. Primary, Adaptive, and Acquired Resistance to Cancer Immunotherapy. Cell 2017; 168: 707–723.

27 Christofi T, Baritaki S, Falzone L, Libra M, Zaravinos A. Current Perspectives in Cancer Immunotherapy. Cancers 2019; 11: 1472.

28 Dunn J, Rao S. Epigenetics and immunotherapy: The current state of play. Mol Immunol 2017; 87: 227–239.

29 Mazzone R, Zwergel C, Mai A, Valente S. Epi-drugs in combination with immunotherapy: a new avenue to improve anticancer efficacy. Clin Epigenetics 2017; 9: 59.

30 Sheng W, LaFleur MW, Nguyen TH, Chen S, Chakravarthy A, Conway JR, Li Y, Chen H, Yang H, Hsu PH, Van Allen EM, Freeman GJ, De Carvalho DD, He HH, Sharpe AH, Shi Y. LSD1 Ablation Stimulates Anti-tumor Immunity and Enables Checkpoint Blockade. Cell 2018; 174: 549–563 e519.

31 Wang J, Hevi S, Kurash JK, Lei H, Gay F, Bajko J, Su H, Sun W, Chang H, Xu G, Gaudet F, Li E, Chen T. The lysine demethylase LSD1 (KDM1) is required for maintenance of global DNA methylation. Nat Genet 2009; 41: 125–129.

32 Tao L, van Bragt MPA, Laudadio E, Li Z. Lineage Tracing of Mammary Epithelial Cells Using Cell-Type-Specific Cre-Expressing Adenoviruses. Stem Cell Reports 2014; 2: 770–779.

33 Tao L, Xiang D, Xie Y, Bronson RT, Li Z. Induced p53 loss in mouse luminal cells causes clonal expansion and development of mammary tumours. Nature communications 2017; 8: 14431.

34 Guy CT, Cardiff RD, Muller WJ. Induction of mammary tumors by expression of polyomavirus middle T oncogene: a transgenic mouse model for metastatic disease. Mol Cell Biol 1992; 12: 954–961.

35 Srinivas S, Watanabe T, Lin CS, William CM, Tanabe Y, Jessell TM, Costantini F. Cre reporter strains produced by targeted insertion of EYFP and ECFP into the ROSA26 locus. BMC Dev Biol 2001; 1: 4.

36 Subramanian A, Tamayo P, Mootha VK, Mukherjee S, Ebert BL, Gillette MA, Paulovich A, Pomeroy SL, Golub TR, Lander ES, Mesirov JP. Gene set enrichment analysis: a knowledge-based approach for interpreting genome-wide expression profiles. Proc Natl Acad Sci U S A 2005; 102: 15545–15550.

37 Treger RS, Tokuyama M, Dong H, Salas-Briceno K, Ross SR, Kong Y, Iwasaki A. Human APOBEC3G Prevents Emergence of Infectious Endogenous Retrovirus in Mice. J Virol 2019; 93.

38 Volkwein W, Pavlovic M, Anton M, Haase M, Stellberger T, Jarrar A, Busch U, Baiker A. Detection and differentiation of murine leukemia virus (MLV) and murine stem cell virus (MSCV) and therefrom derived nucleic acids. J Virol Methods 2022; 299: 114316.

39 Krneta T, Gillgrass A, Chew M, Ashkar AA. The breast tumor microenvironment alters the phenotype and function of natural killer cells. Cell Mol Immunol 2016; 13: 628–639.

40 Andre P, Denis C, Soulas C, Bourbon-Caillet C, Lopez J, Arnoux T, Blery M, Bonnafous C, Gauthier L, Morel A, Rossi B, Remark R, Breso V, Bonnet E, Habif G, Guia S, Lalanne AI, Hoffmann C, Lantz O, Fayette J, Boyer-Chammard A, Zerbib R, Dodion P, Ghadially H, Jure-Kunkel M, Morel Y, Herbst R, Narni-Mancinelli E, Cohen RB, Vivier E. Anti-NKG2A mAb Is a Checkpoint Inhibitor that Promotes Anti-tumor Immunity by Unleashing Both T and NK Cells. Cell 2018; 175: 1731–1743 e1713.

41 Cao X, Shores EW, Hu-Li J, Anver MR, Kelsall BL, Russell SM, Drago J, Noguchi M, Grinberg A, Bloom ET, et al. Defective lymphoid development in mice lacking expression of the common cytokine receptor gamma chain. Immunity 1995; 2: 223–238.

42 Fung-Leung WP, Schilham MW, Rahemtulla A, Kundig TM, Vollenweider M, Potter J, van Ewijk W, Mak TW. CD8 is needed for development of cytotoxic T cells but not helper T cells. Cell 1991; 65: 443–449.

43 Lopez-Soto A, Gonzalez S, Smyth MJ, Galluzzi L. Control of Metastasis by NK Cells. Cancer Cell 2017; 32: 135–154.

44 Long EO, Kim HS, Liu D, Peterson ME, Rajagopalan S. Controlling natural killer cell responses: integration of signals for activation and inhibition. Annu Rev Immunol 2013; 31: 227–258.

45 He Y, Tian Z. NK cell education via nonclassical MHC and non-MHC ligands. Cell Mol Immunol 2017; 14: 321–330.

46 Muntasell A, Ochoa MC, Cordeiro L, Berraondo P, Lopez-Diaz de Cerio A, Cabo M, Lopez-Botet M, Melero I. Targeting NK-cell checkpoints for cancer immunotherapy. Curr Opin Immunol 2017; 45: 73–81.

47 Rautela J, Baschuk N, Slaney CY, Jayatilleke KM, Xiao K, Bidwell BN, Lucas EC, Hawkins ED, Lock P, Wong CS, Chen W, Anderson RL, Hertzog PJ, Andrews DM, Moller A, Parker BS. Loss of Host Type-I IFN Signaling Accelerates Metastasis and Impairs NK-cell Antitumor Function in Multiple Models of Breast Cancer. Cancer Immunol Res 2015; 3: 1207–1217.

48 Zeng L, Sullivan LC, Vivian JP, Walpole NG, Harpur CM, Rossjohn J, Clements CS, Brooks AG. A structural basis for antigen presentation by the MHC class Ib molecule, Qa-1b. Journal of immunology 2012; 188: 302–310.

49 Jensen PE, Sullivan BA, Reed-Loisel LM, Weber DA. Qa-1, a nonclassical class I histocompatibility molecule with roles in innate and adaptive immunity. Immunol Res 2004; 29: 81–92.

50 Braud V, Jones EY, McMichael A. The human major histocompatibility complex class Ib molecule HLA-E binds signal sequence-derived peptides with primary anchor residues at positions 2 and 9. European journal of immunology 1997; 27: 1164–1169.

51 Braud VM, Allan DS, Wilson D, McMichael AJ. TAP- and tapasin-dependent HLA-E surface expression correlates with the binding of an MHC class I leader peptide. Curr Biol 1998; 8: 1–10.

52 Iwaszko M, Bogunia-Kubik K. Clinical significance of the HLA-E and CD94/NKG2 interaction. Arch Immunol Ther Exp (Warsz) 2011; 59: 353–367.

53 Vance RE, Kraft JR, Altman JD, Jensen PE, Raulet DH. Mouse CD94/NKG2A is a natural killer cell receptor for the nonclassical major histocompatibility complex (MHC) class I molecule Qa-1(b). J Exp Med 1998; 188: 1841–1848.

54 Braud VM, Allan DS, O’Callaghan CA, Soderstrom K, D’Andrea A, Ogg GS, Lazetic S, Young NT, Bell JI, Phillips JH, Lanier LL, McMichael AJ. HLA-E binds to natural killer cell receptors CD94/NKG2A, B and C. Nature 1998; 391: 795–799.

55 Shultz LD, Schweitzer PA, Christianson SW, Gott B, Schweitzer IB, Tennent B, McKenna S, Mobraaten L, Rajan TV, Greiner DL, et al. Multiple defects in innate and adaptive immunologic function in NOD/LtSz-scid mice. Journal of immunology 1995; 154: 180–191.

56 DeNardo DG, Barreto JB, Andreu P, Vasquez L, Tawfik D, Kolhatkar N, Coussens LM. CD4(+) T cells regulate pulmonary metastasis of mammary carcinomas by enhancing protumor properties of macrophages. Cancer Cell 2009; 16: 91–102.

57 Lin EY, Nguyen AV, Russell RG, Pollard JW. Colony-stimulating factor 1 promotes progression of mammary tumors to malignancy. J Exp Med 2001; 193: 727–740.

58 Curtis C, Shah SP, Chin SF, Turashvili G, Rueda OM, Dunning MJ, Speed D, Lynch AG, Samarajiwa S, Yuan Y, Graf S, Ha G, Haffari G, Bashashati A, Russell R, McKinney S, Langerod A, Green A, Provenzano E, Wishart G, Pinder S, Watson P, Markowetz F, Murphy L, Ellis I, Purushotham A, Borresen-Dale AL, Brenton JD, Tavare S, Caldas C, Aparicio S. The genomic and transcriptomic architecture of 2,000 breast tumours reveals novel subgroups. Nature 2012; 486: 346–352.

59 Pereira B, Chin SF, Rueda OM, Vollan HK, Provenzano E, Bardwell HA, Pugh M, Jones L, Russell R, Sammut SJ, Tsui DW, Liu B, Dawson SJ, Abraham J, Northen H, Peden JF, Mukherjee A, Turashvili G, Green AR, McKinney S, Oloumi A, Shah S, Rosenfeld N, Murphy L, Bentley DR, Ellis IO, Purushotham A, Pinder SE, Borresen-Dale AL, Earl HM, Pharoah PD, Ross MT, Aparicio S, Caldas C. The somatic mutation profiles of 2,433 breast cancers refines their genomic and transcriptomic landscapes. Nature communications 2016; 7: 11479.

60 Qin Y, Vasilatos SN, Chen L, Wu H, Cao Z, Fu Y, Huang M, Vlad AM, Lu B, Oesterreich S, Davidson NE, Huang Y. Inhibition of histone lysine-specific demethylase 1 elicits breast tumor immunity and enhances antitumor efficacy of immune checkpoint blockade. Oncogene 2019; 38: 390–405.

61 Bailey CP, Figueroa M, Gangadharan A, Lee DA, Chandra J. Scaffolding LSD1 Inhibitors Impair NK Cell Metabolism and Cytotoxic Function Through Depletion of Glutathione. Front Immunol 2020; 11: 2196.

62 Xiang D, Tao L, Li Z. Modeling Breast Cancer via an Intraductal Injection of Cre-expressing Adenovirus into the Mouse Mammary Gland. J Vis Exp 2019; (148): 10.3791/59502.

63 Van Keymeulen A, Rocha AS, Ousset M, Beck B, Bouvencourt G, Rock J, Sharma N, Dekoninck S, Blanpain C. Distinct stem cells contribute to mammary gland development and maintenance. Nature 2011; 479: 189–193.

64 Shackleton M, Vaillant F, Simpson KJ, Stingl J, Smyth GK, Asselin-Labat ML, Wu L, Lindeman GJ, Visvader JE. Generation of a functional mammary gland from a single stem cell. Nature 2006; 439: 84–88.

65 Wang H, Xiang D, Liu B, He A, Randle HJ, Zhang KX, Dongre A, Sachs N, Clark AP, Tao L, Chen Q, Botchkarev VV, Jr., Xie Y, Dai N, Clevers H, Li Z, Livingston DM. Inadequate DNA Damage Repair Promotes Mammary Transdifferentiation, Leading to BRCA1 Breast Cancer. Cell 2019; 178: 135–151 e119.

66 Neri S, Mariani E, Meneghetti A, Cattini L, Facchini A. Calcein-acetyoxymethyl cytotoxicity assay: standardization of a method allowing additional analyses on recovered effector cells and supernatants. Clin Diagn Lab Immunol 2001; 8: 1131–1135.

67 Ohtsuka M, Inoko H, Kulski JK, Yoshimura S. Major histocompatibility complex (Mhc) class Ib gene duplications, organization and expression patterns in mouse strain C57BL/6. BMC Genomics 2008; 9: 178.

68 Shalem O, Sanjana NE, Hartenian E, Shi X, Scott DA, Mikkelson T, Heckl D, Ebert BL, Root DE, Doench JG, Zhang F. Genome-scale CRISPR-Cas9 knockout screening in human cells. Science 2014; 343: 84–87.

69 Follenzi A, Ailles LE, Bakovic S, Geuna M, Naldini L. Gene transfer by lentiviral vectors is limited by nuclear translocation and rescued by HIV-1 pol sequences. Nat Genet 2000; 25: 217–222.

70 Schwank G, Koo BK, Sasselli V, Dekkers JF, Heo I, Demircan T, Sasaki N, Boymans S, Cuppen E, van der Ent CK, Nieuwenhuis EE, Beekman JM, Clevers H. Functional repair of CFTR by CRISPR/Cas9 in intestinal stem cell organoids of cystic fibrosis patients. Cell Stem Cell 2013; 13: 653–658.

