## Supplementary Figures S1-S6 for "LSD1 ablation promotes mammary tumor metastasis by attenuating NK cell-mediated anti-tumor immunity"

### Supplementary figures and figure legends

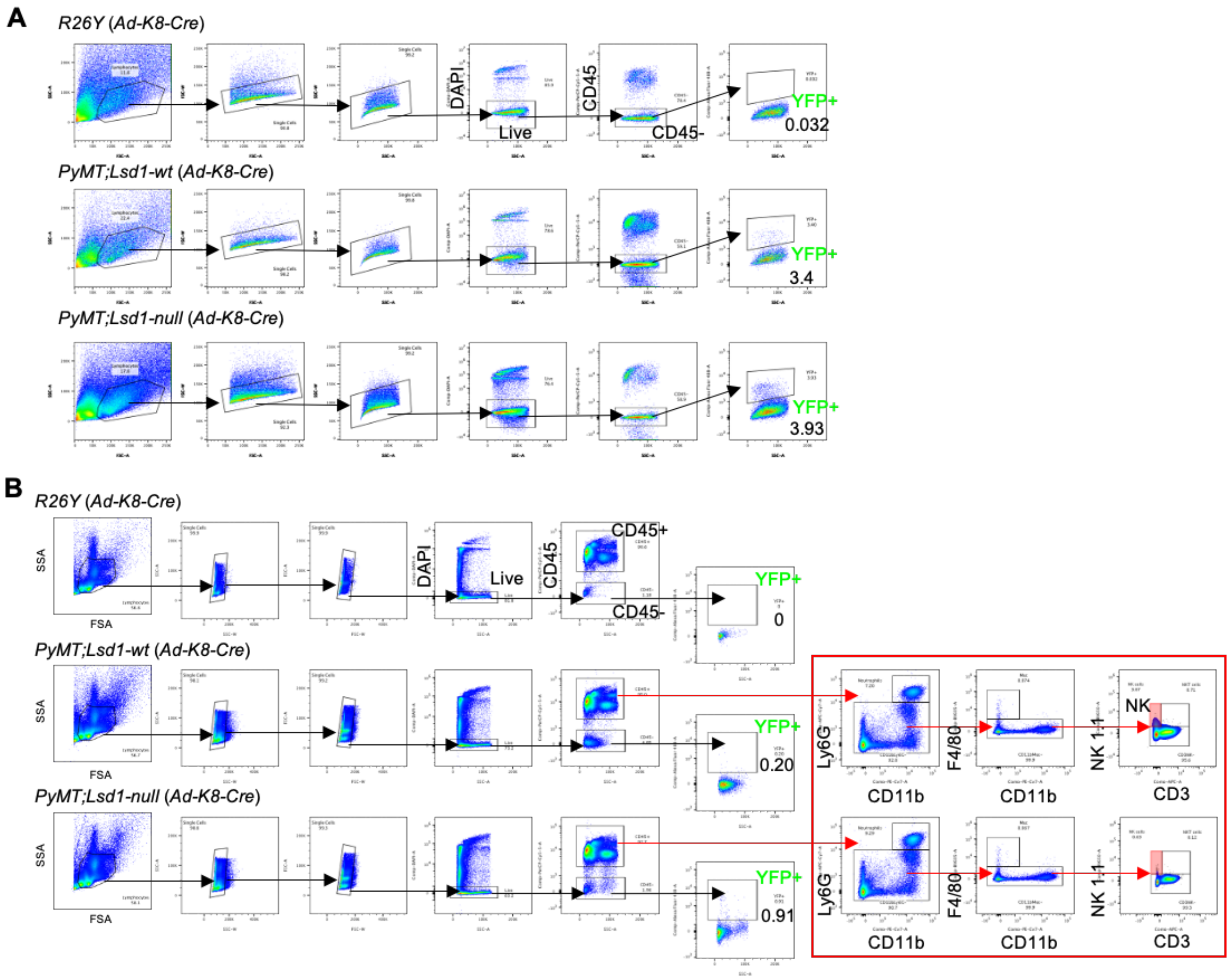


**Figure S1. Analysis of YFP⁺ cells and NK cells. (A)** Representative FACS gating for YFP⁺ cells from CD45⁻ cells in *PyMT* primary mammary tumors. **(B)** FACS gating showing the proportion of circulating YFP⁺ tumor cells in the peripheral blood samples from *PyMT* mice with or without induced LSD1-loss, as well as NK cells among CD45⁺ cells in the peripheral blood samples. In both (A) and (B), wild-type female mice with the *R26Y* reporter alone were used as a control to set up the FACS gates (upon intraductal injection of *Ad-K8-Cre*). Both *PyMT;Lsd1-wt* and *PyMT;Lsd1-null* mice here also carried the same *R26Y* reporter.

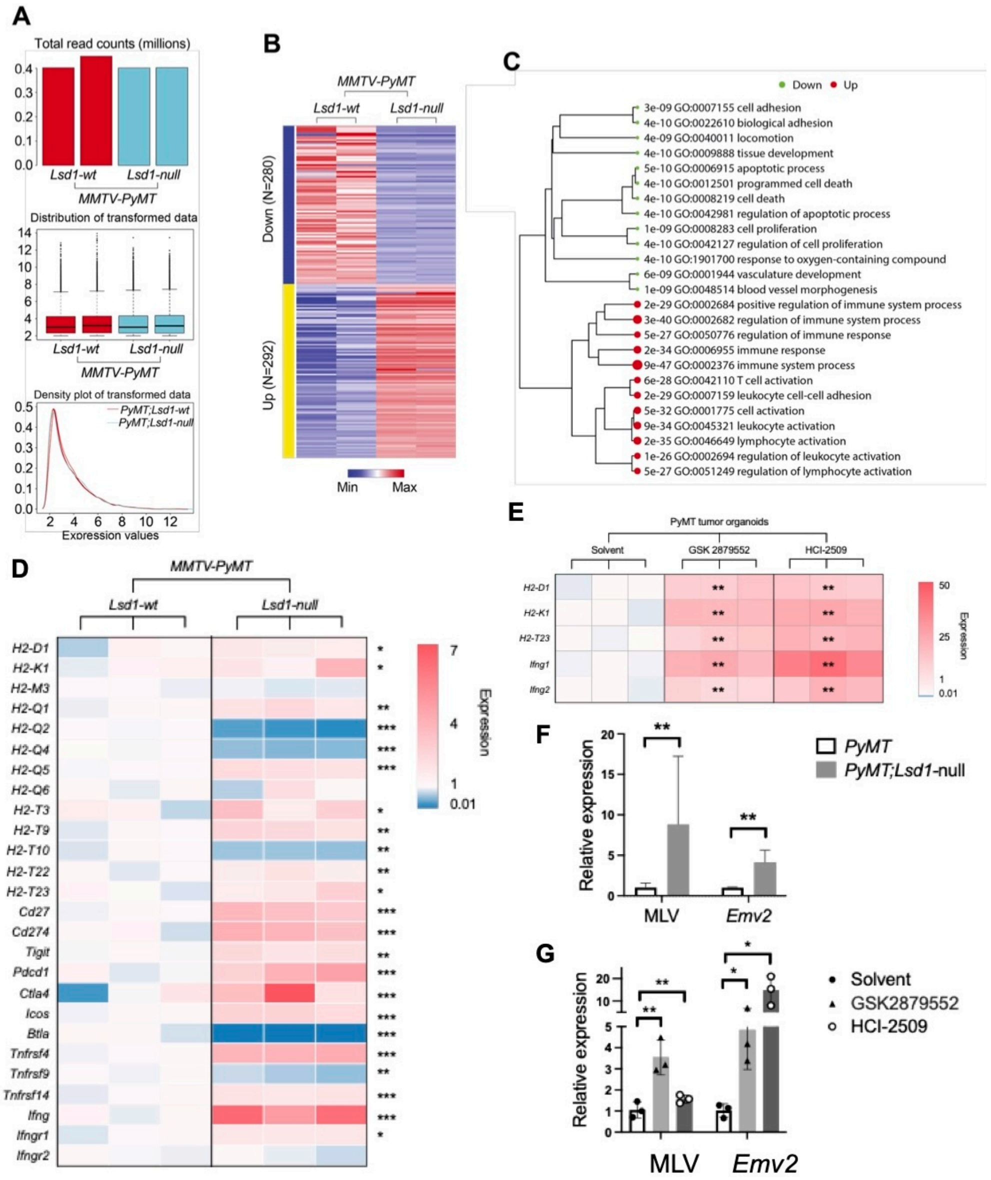


**Figure S2. Expression analysis. (A-C)** Graphical summary of Viper analysis for YFP+ *PyMT* tumor cells with/without LSD1-loss: **(A)** Total read counts of indicated samples. **(B)** Heatmap of transformed read counts for genes with adjusted p-values <0.05 in the DGE analysis. **(C)** Dot plot generated with the clusterProfiler package showing significantly enriched pathways. **(D)** Heatmaps showing the expression of classical and non-classical MHC-I genes, as well as IFNγ(*Ifng*) and its receptor genes in *PyMT* tumor organoids with scramble control (*Lsd1*-wt) and CRIPSR-mediated *Lsd1* knockout (*Lsd1*-null). **(E)** Differential expression of selected non-MHC-I genes and IFNγ receptor genes in *PyMT* tumor organoids with or without different LSD1 inhibitor treatment (GSK2879552 or HCI-2509). **(F-G)** qRT-PCR data for expression of ERV genes in sorted YFP+ *Lsd1*-null versus wildtype *PyMT* tumor cells (F) and *PyMT* tumor organoids upon different LSD1 inhibitor treatment (G). *P* value: *p<0.05, **p<0.01, ***p<0.005, two-tailed Student's *t*-test. Data represent mean ± SEM.

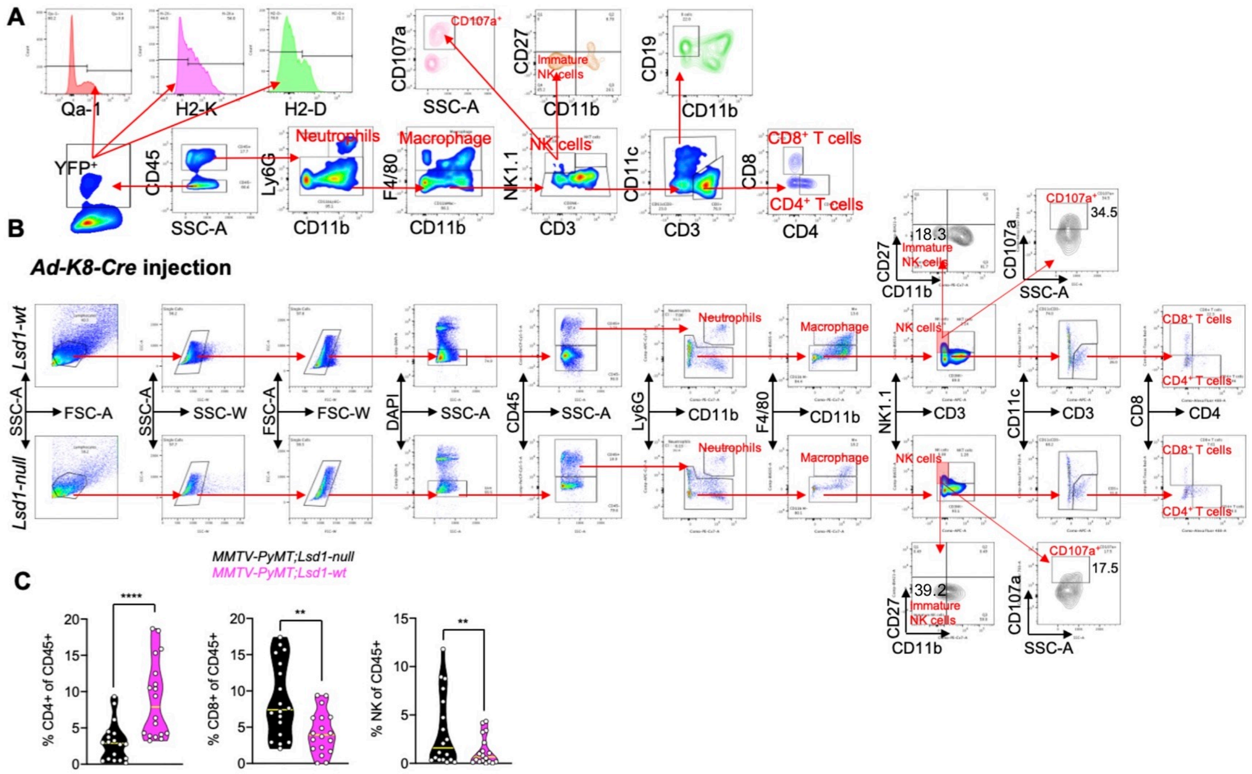


**Figure S3. FACS gating strategy for live CD45$^{+}$ immune cells in the *PyMT* tumor microenvironment (*Ad-K8-Cre*-based model). (A)** FACS gating strategy for live CD45$^{+}$ immune cells in the tumor microenvironment. **(B)** FACS gating strategy for various live CD45$^{+}$ immune cell subsets from *PyMT* tumors with *Ad-K8-Cre* injection. **(C)** Alteration of the representative immune cell subsets in *PyMT* tumors with LSD1-loss (*Lsd1-null*), based on the FACS analysis shown in (B). *P* value: **p<0.01, ****p<0.001, two-tailed Student's t-test. Data represent mean ± SEM.

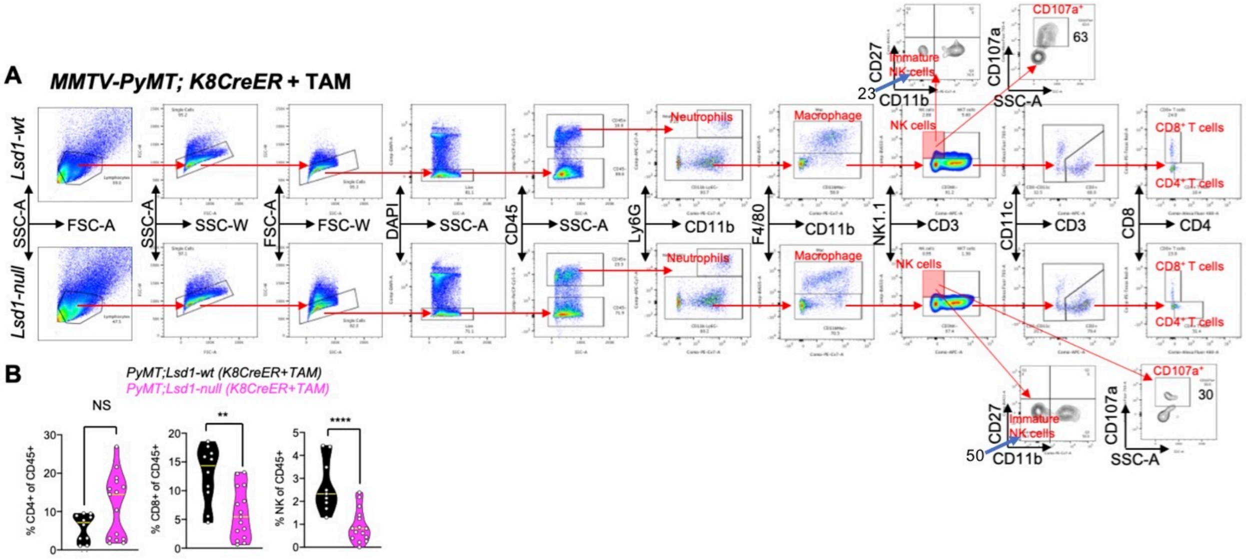


**Figure S4. FACS gating strategy for live CD45$^+$ immune cells in the *PyMT* tumor microenvironment (*K8-CreER*-based model). (A)** FACS gating strategy for live CD45$^+$ immune cells from *PyMT* tumors with tamoxifen (TAM) induction of K8-CreER. **(B)** Alteration of the representative immune cell subsets in *PyMT* tumors with TAM induction of K8-CreER, based on the FACS analysis shown in (A).

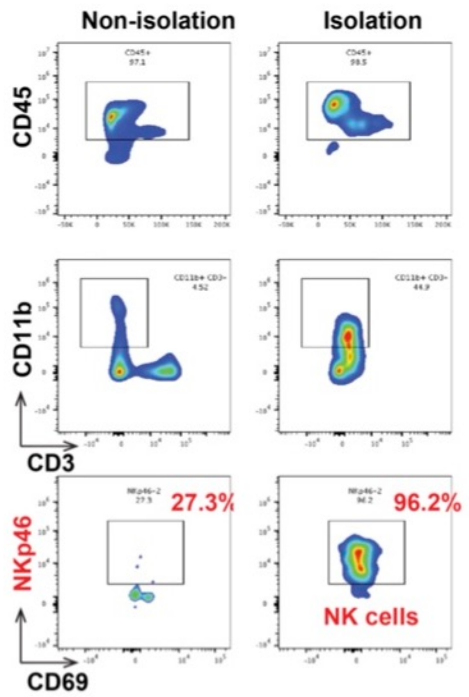


**Figure S5. Isolation of NK cells.** Typical mouse NK Cell isolation profile. Starting with mouse bone marrow cells, the purities of the start and enriched NK cell ($CD45^{+}CD3^{-}CD69^{+}NKp46^{+}$) fractions were 27.3% and 96.2%, respectively.

**A**

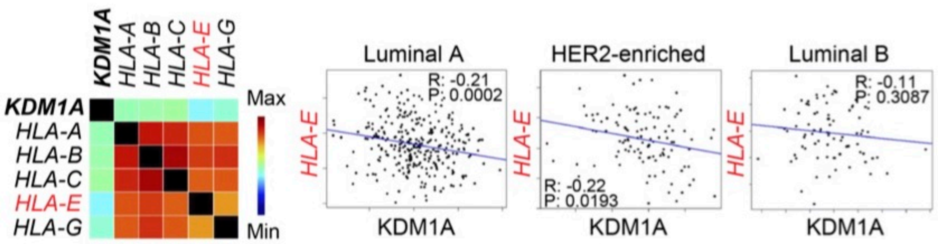


**C**

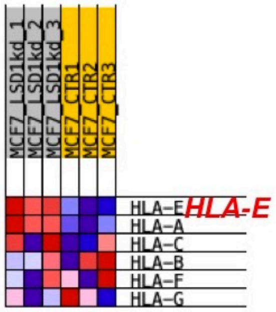


**B**

| RANK | NAME | SIZE | ES | NES | NOM p-val | FDR q-val |
|---|---|---|---|---|---|---|
| 1 | WP_TYROBP_CAUSAL_NETWORK | 57 | 0.72 | 2.25 | 0 | 0 |
| 2 | REACTOME_IMMUNOREGULATORY_INTERACTIONS_BETWEEN_A_LYMPHOID_AND_A_NON_LYMPHOID_CELL | 108 | 0.63 | 2.19 | 0 | 0 |
| 3 | KEGG_RETINOL_METABOLISM | 50 | 0.7 | 2.17 | 0 | 0 |
| 4 | KEGG_INTESTINAL_IMMUNE_NETWORK_FOR_IGA_PRODUCTION | 45 | 0.7 | 2.12 | 0 | 2.79E-04 |
| 5 | KEGG_SYSTEMIC_LUPUS_ERYTHEMATOSUS | 123 | 0.6 | 2.12 | 0 | 2.23E-04 |
| 6 | REACTOME_INTERLEUKIN_10_SIGNALING | 45 | 0.69 | 2.11 | 0 | 3.69E-04 |
| 7 | REACTOME_CHEMOKINE_RECEPTORS_BIND_CHEMOKINES | 53 | 0.68 | 2.11 | 0 | 3.17E-04 |
| 8 | WP_ALLOGRAFT_REJECTION | 87 | 0.63 | 2.1 | 0 | 4.15E-04 |
| 9 | WP_HUMAN_COMPLEMENT_SYSTEM | 88 | 0.63 | 2.09 | 0 | 4.92E-04 |
| 10 | KEGG_GRAFT_VERSUS_HOST_DISEASE | 36 | 0.73 | 2.08 | 0 | 7.75E-04 |
| 11 | REACTOME_NEUTROPHIL_DEGRANULATION | 444 | 0.52 | 2.07 | 0 | 7.05E-04 |
| 12 | WP_SPINAL_CORD_INJURY | 115 | 0.59 | 2.06 | 0 | 7.35E-04 |
| 13 | KEGG_PPAR_SIGNALING_PATHWAY | 63 | 0.64 | 2.05 | 0 | 6.79E-04 |
| 14 | REACTOME_SYNTHESIS_SECRETION_AND_DEACYLATION_OF_GHRELIN | 17 | 0.84 | 2.05 | 0 | 6.30E-04 |
| 15 | WP_PPAR_SIGNALING_PATHWAY | 62 | 0.64 | 2.05 | 0 | 5.88E-04 |
| 16 | KEGG_AUTOIMMUNE_THYROID_DISEASE | 48 | 0.67 | 2.04 | 0 | 5.52E-04 |
| 17 | BIOCARTA_COMP_PATHWAY | 18 | 0.8 | 2.04 | 0 | 7.76E-04 |
| 18 | REACTOME_COMPLEMENT_CASCADE | 51 | 0.66 | 2.03 | 0 | 0.00109896 |
| 19 | REACTOME_INITIAL_TRIGGERING_OF_COMPLEMENT | 21 | 0.78 | 2.03 | 0 | 0.00104112 |
| 20 | REACTOME_TRIGLYCERIDE_METABOLISM | 35 | 0.71 | 2.03 | 0 | 9.89E-04 |
| 21 | KEGG_ALLOGRAFT_REJECTION | 35 | 0.71 | 2.02 | 0 | 0.00104621 |
| 22 | KEGG_TYPE_I_DIABETES_MELLITUS | 40 | 0.68 | 2.02 | 0 | 0.00110183 |
| 23 | REACTOME_ANTIGEN_PROCESSING_CROSS_PRESENTATION | 91 | 0.6 | 2.01 | 0 | 0.00129212 |
| 24 | KEGG_HEMATOPOIETIC_CELL_LINEAGE | 76 | 0.61 | 2 | 0 | 0.00146641 |
| 25 | WP_COMPLEMENT_ACTIVATION | 22 | 0.76 | 1.99 | 0 | 0.0016742 |
| 26 | KEGG_ASTHMA | 28 | 0.73 | 1.98 | 0 | 0.00178032 |
| 27 | REACTOME_INNATE_IMMUNE_SYSTEM | 963 | 0.47 | 1.98 | 0 | 0.00216498 |
| 28 | KEGG_LEISHMANIA_INFECTION | 62 | 0.63 | 1.97 | 0 | 0.00216699 |
| 29 | WP_ELECTRON_TRANSPORT_CHAIN_OXPHOS_SYSTEM_IN_MITOCHONDRIA | 81 | 0.58 | 1.96 | 0 | 0.00270049 |
| 30 | WP_PROSTAGLANDIN_SYNTHESIS_AND_REGULATION | 42 | 0.66 | 1.96 | 0 | 0.00261047 |

**Figure S6. Immune signature in *LSD1*-low human luminal breast cancer cells. (A)** Left: heatmap showing correlation of expression of *LSD1* (*KDM1A*) with several *HLA* genes in human luminal A breast cancers; right: correlation of *HLA-E* and *KDM1A* expression in different subtypes of human breast cancers. The analysis was based on the Breast Cancer Gene-Expression Miner (bc-GenExMiner) online tool. **(B)** GSEA showing top enriched gene sets (from the GSEA MSigDB C2 canonical pathways collection) in *LSD1*-low group (in relation to *LSD1*-high group, as in Figure 7A-B); immune-related gene sets are highlighted in red. ES: enrichment score; NES: normalized enrichment score; NOM p-val: nominal p-value; FDR q-val: false discovery rate q-value. **(C)** Heatmap (generated using GSEA and *HLA* genes as a gene set; red to blue indicate highest to lowest expression) showing upregulation of *HLA-E* in MCF7 cells with *LSD1* (*KDM1A*) knockdown, the data is based on GEO accession #: GSE95165.
